# Acidic transcription factors position the genome at nuclear speckles through transcription dependent and independent mechanisms

**DOI:** 10.1101/2025.10.01.679912

**Authors:** Pankaj Chaturvedi, Purnam Ghosh, Liguo Zhang, Meng Zhang, Huimin Zhao, Andrew S. Belmont

## Abstract

A small fraction of the genome positions reproducibly near nuclear speckles (NS), increasing expression and/or splicing efficiency of NS-associated genes. How specific genomic regions target to NS remains unclear. Here, we demonstrate that establishment of genome-wide NS-association occurs independent of active transcription. We show that DNA sequences derived from NS-associated regions integrated as transgenes autonomously target to NS. By systematically dissecting one such genomic locus, the COL1A1-SGCA locus, we identified redundant NS-targeting cis regulatory elements, including a ∼600 bp fragment with 17 binding motifs for 8 transcription factors (TFs). Four NS-targeting TFs within this fragment contain acidic activation domains (AADs) that provide both chromatin-context and transcription-dependent NS-targeting, a property that appears common among several other tested AADs. A subset of acidic activator TFs contain an additional, transcription-independent NS-targeting activity. Our findings establish diverse and partially redundant NS-targeting activities, which may facilitate dynamic gene positioning at NS periphery for context-specific transcriptional responses.

## INTRODUCTION

The spatial organization of metazoan nuclei into distinct membraneless subcompartments increases the local concentration of regulatory macromolecules, thereby influencing the distribution, dynamics, and interactions of the segregated factors. Nuclear speckles (NS) represent one such nuclear subcompartment. NS are enriched in factors related to multiple stages of RNA Pol II gene expression, including different steps of transcription, RNA processing, and RNA export.^1–5^ NS are irregularly shaped with variable size (∼0.3-3.0 μm) and numbers (∼20-50) per nucleus.^6–9^ Depending on the methods of detection, NS have also been referred to as hyaline grumes,^10^ interchromatin granule clusters (IGCs),^7,11^ poly(A) RNA domains,^12^ SC-35 domains,^6^ splicing factor compartments (SFCs),^13^ or B snurposomes in amphibian oocytes.^11^ Even the definition of the NS periphery varies with the NS markers used for their detection.^5^ SON and SRRM2 are considered to be core NS proteins.^14,15^ To operationally define NS, we used SON immunostaining which appears to closely correspond to the light microscopy equivalent of IGCs.

*In situ* hybridization of >25 actively transcribed genes had earlier demonstrated close NS-association of more than half of these genes.^8,14,16^ The nascent RNAs from two of these NS-associated genes (COL1A1 and cardiac myosin heavy chain (cMyHC)) entered the adjacent NS, whereas the nascent RNAs of the other NS-associated genes (β-actin (*ACTB*), transcription factor E2F4, lamin A/C (*LMNA*), and fibronectin (*FN1*)) localized outside the adjacent NS.^16,17^ Therefore, NS-associated genes were subdivided into Type 1 genes, whose transcripts accumulated within NS, versus Type 2 genes, whose transcripts accumulated next to the gene without entering the NS.^16,17^ These observations led to a “gene expression hub” model in which NS were suggested as constituting a special nuclear compartment facilitating the expression of a subset of genes.^14^

More recently, genome-wide sequencing (e.g. TSA-Seq and SPRITE)^18–20^ and imaging^21,22^ approaches confirm preferential positioning of a significant fraction of active chromosomal regions near NS. For example, TSA-Seq identified that approximately half of the top 5% highly expressed genes are positioned within several hundred nanometer (nm) from the NS periphery.^18^ This enrichment of the most highly expressed genes near NS was consistent across four tested cell lines.^20^ Additionally, proximity to NS has been shown to increase gene expression either through overall amplified expression,^5,23,24^ and/or through enhanced RNA splicing efficiency.^25,26^

A recent study highlights the role of the NS periphery in amplifying gene expression by showing the downregulation of 100s of NS-proximal genes after elimination of NS by SON and SRRM2 double knockdown.^27^ Therefore, targeting to NS periphery appears to be an important determinant for the expression of such genes although the underlying mechanisms for NS-targeting of these genes are largely unknown. Previously, we demonstrated the autonomous NS-targeting of HSPA1 BAC transgenes to NS,^28^ which closely reproduced the NS-targeting of the endogenous *HSPA1* locus. This NS-targeting by HSPA1 BAC transgene was significantly reduced after transcriptional inhibition.^28^ In contrast, earlier studies by Lawrence group showed that the NS-associated gene COL1A1 associates stably with NS even after transcriptional inhibition.^17^ These results suggested the possibility of at least two different NS-targeting mechanisms.

Here we used a top-down approach to identify first cis and then trans elements conferring either transcription-dependent or transcription-independent NS-targeting. Using TSA-Seq, we first showed that the establishment and maintenance of the overall positioning of endogenous chromosomes relative to NS is transcription-independent.

We next demonstrated the autonomous, integration-site independent targeting to NS of multiple 100-200 kbp DNA sequences derived from NS-associated domains (SPADs) after random genome integration. Using this autonomous targeting activity of transgenes as our assay, we identified multiple functionally redundant cis regulatory elements (CREs) conferring either transcription-dependent or -independent NS-targeting. This dissection then led us to the demonstration of transcription-dependent NS-targeting activity as a common property of acidic transcription factors (TF), with a subset of these acidic TFs showing both transcription-dependent and -independent NS-targeting activities localizing to different TF domains.

Our results suggest a model for mammalian genome organization where the combinatorial action of multiple, functionally redundant CREs containing binding motifs for acidic TFs provide targeting and stable anchoring of a subset of chromosome regions to the NS periphery.

## RESULTS

### Genome organization relative to NS is maintained and established independent of transcription

To study the effect of transcription on genome-NS-association, we applied two commonly used transcriptional inhibitors, Triptolide (TPL) and 5,6-Dichloro-1-β-D-ribofuranosylbenzimidazole (DRB). TPL irreversibly inhibits XPB/TFIIH at transcriptional initiation and induces rapid degradation of RNA Pol II,^29^ while DRB targets the CDK9 kinase subunit of P-TEFβ and reversibly inhibits transcriptional elongation.^30^ TPL treatment at 125 nM concentration for 1 hour itself is sufficient to induce global transcription block, however ∼1% genes still retain paused Pol II on their promoters.^29^ Therefore, to achieve maximum transcriptional inhibition in asynchronously proliferating hTERT-immortalized RPE1 we performed independent transcriptional inhibition experiments for extended periods (4 hours) with TPL or DRB. Cells treated with the inhibitors showed a dramatic reduction in both nascent transcription, assayed by 5-Ethynyl uridine (EU, a ribonucleotide homolog) pulse-labeling (Figure 1A, left panel, Figure 1B), and RNA Pol II -Ser5 phosphorylation, assayed by immunostaining (Figure 1A, middle panel, Figure 1C). As previously reported,^3,31^ treatment of cells with these inhibitors also resulted in a rounding of NS. After the 4-hour DRB treatment, a small number of speckles in ∼40% of cells showed dramatic enlargement (Figure 1A), probably due to the fusion of speckles, as reported in our previous study.^31^

**Figure 1.**
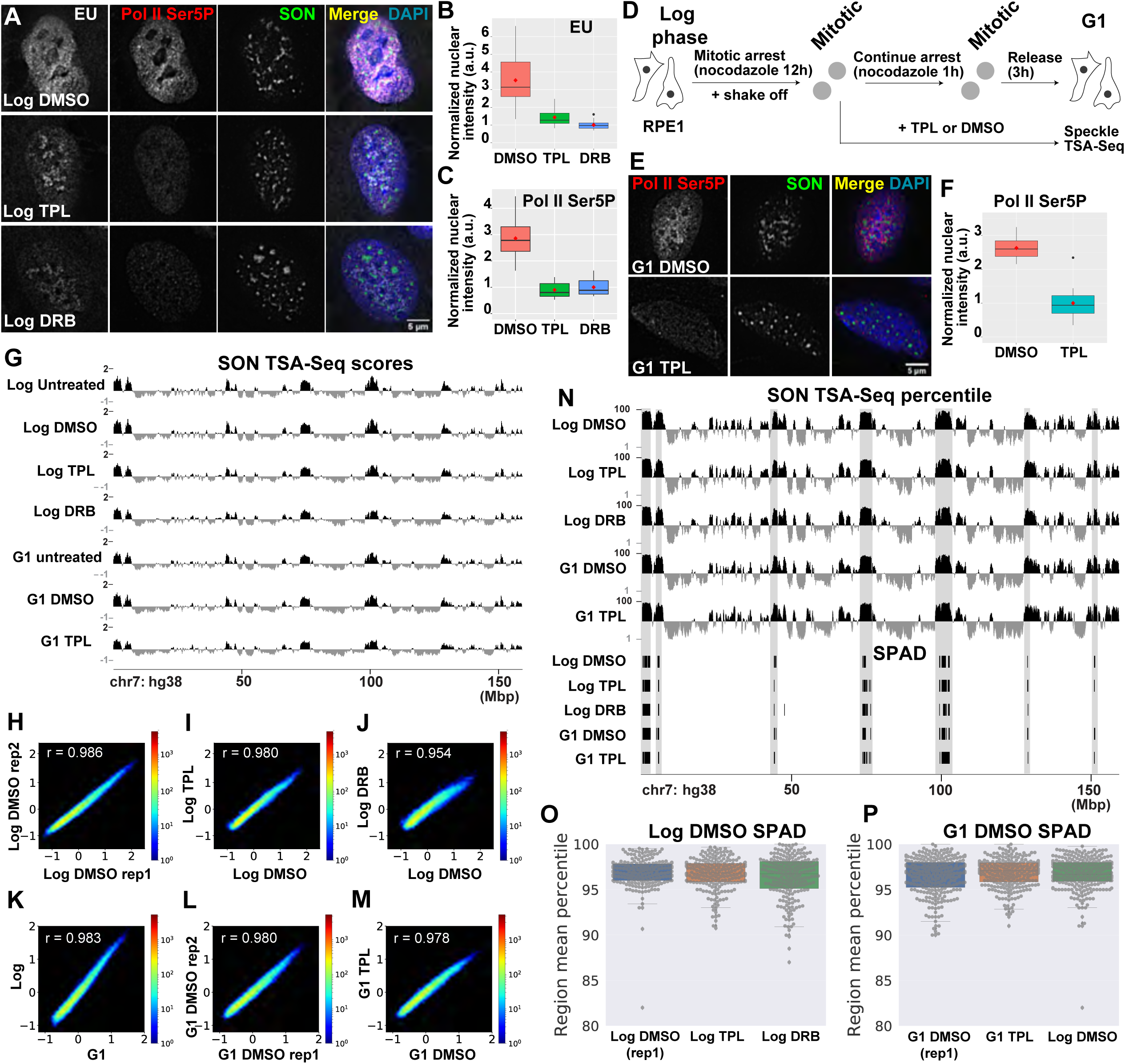
Genome organization relative to NS is maintained and established independent of transcription. **(A-C)** Validation of transcription inhibition **(A)** EU pulse label, Pol II Ser5P, and SON immunostaining merged with DAPI in RPE1 cells treated with DMSO (top), TPL (middle), or DRB (bottom). **(B-C)** Quantification EU and Pol II Ser5p staining in (A) Intensities were normalized to the mean of the DRB samples. Box plots show mean (red dots), median (black horizontal line), interquartile range (IQR) (box), range within1.5x IQR (top and bottom whiskers), and individual data points. n = 20-23 nuclei. **(D)** Schematic showing RPE1 cell synchronization and transcription inhibition. **(E)** Comparison of Pol II Ser5P and SON immunostaining with DAPI merge in G1 phase RPE1 cells treated beginning in mitosis with DMSO (top) or TPL (bottom). **(F)** Nuclear Pol II Ser5P intensities of staining in (E) normalized to the TPL sample mean. n = 14 cells. **(G-O)** Minimal perturbation of G1 genome-wide NS association patterns after TPL inhibition beginning in mitosis (G1 samples) or log phase. (G) Genome browser view of SON TSA-Seq enrichment scores in untreated, DMSO-, TPL-, or DRB-treated log phase or G1 phase RPE1 cells. **(H-M)** Two-dimensional histograms comparing SON TSA-Seq enrichment scores between replicate DMSO-treated samples (**H**: log phase, **L**: G1 phase) or between DMSO and TPL (**I**: log phase, **M**: G1 phase) or between DMSO and DRB treatment (**J**: log phase) or between untreated log versus G1 phase cells (**K**). Colors indicate the number of 20 kb genomic bins within each 2D TSA-Seq score interval (∼0.02 x 0.02). Pearson’s r indicates the Pearson product-moment correlation coefficient. **(N)** SON TSA-Seq score percentile tracks (20 kb bins, top), and segmented Speckle-Associated Domains (SPADs, bottom) in log phase RPE1 cells treated with DMSO, TPL, or DRB, or in G1 phase cells treated with DMSO or TPL. **(O–P)** Distributions of SON TSA-Seq percentiles for SPADs from DMSO-treated log phase **(O)** or G1 phase **(P)** cells compared with biological replicates, TPL-treated, or DRB-treated cells. Box plots show the median (black line), 25th–75th percentiles (box), whiskers (within 1.5× IQR), outliers (diamonds), and individual SPAD values (gray dots). Image panel scale bars = 5 μm.

We applied the TSA-Seq 2.0 method for mapping genome organization relative to NS using SON as the marker protein in asynchronously proliferating RPE1 cells, or to cells arrested in mitosis and then synchronously released into G1 phase (Figure 1D).^15,18,20^ Similar to our observations with asynchronous RPE1 cells, the level of RNA Pol II-Ser5 phosphorylation was also reduced in synchronized G1 phase cells upon TPL treatment (Figure 1E-F).

Overall NS TSA-Seq profiles remained largely unchanged after treatment of both asynchronous or G1 synchronized RPE1 cells with transcriptional inhibitors as compared to untreated cells or the DMSO control (Figure 1G). A strong correlation was observed between transcriptionally-inhibited cells and control cells (0.954 – 0.980, Pearson’s *r*), and between biological replicates of the DMSO control cells (*r* = 0.986) (Figure 1H-M). A similar conservation of overall NS TSA-Seq patterns comparing control versus TPL treated samples was observed in a different cell line-asynchronously proliferating HCT116 cells (Figure S1).

We therefore conclude that maintenance of NS-association of speckle associated domains (SPADs) does not require ongoing transcription.

While the above results demonstrated that ongoing transcription was not required to maintain NS-association of SPADs over several hours, they did not rule out the possibility that transcription was required to establish NS-association of SPADs. To address this question, we blocked cells in mitosis, inhibited transcription and released the cells into G1 with a continued inhibition of transcription during which time the NS reformed in these early G1 cells. We performed SON TSA-Seq 3 hours after release of mitotic block. By this time cells had exited mitosis and reformed interphase nuclei with decondensed chromosomes and typical NS morphology.

To gather the large numbers of cells required for TSA-Seq, we first accumulated large numbers of log phase hTERT-RPE1 cells in mitosis using a 12-hour nocodazole treatment (Figure 1D). We then collected mitotic cells using a mechanical shake-off and incubated these cells with TPL for one hour to ensure transcriptional inhibition while still holding them in mitosis with continued nocodazole treatment. We then released cells into G1 by removing the nocodazole while maintaining TPL treatment for another 3 hours (Figure 1D). After this 3-hour release from nocodazole-block, >95% of all mitotically arrested cells were able to enter G1 phase, although entry was slightly delayed for TPL treated cells compared to untreated or DMSO control cells, as measured by DAPI-staining of cells at different time points during this 3-hour period.

The level of transcriptional inhibition in these G1 cells was comparable to that achieved in log phase cells, as evaluated by RNA Pol II Ser5P immunostaining (Figure 1C&F).

We then performed a genome-wide mapping relative to NS using SON TSA-Seq 2.0 after this 3-hour release from the nocodazole block with or without transcriptional inhibition by TPL. Again, we observed very similar NS TSA-Seq profiles in the transcription-inhibited versus control G1 cells, which were both similar to the NS TSA-Seq profiles from asynchronously proliferating cells (Figure 1G). Pearson’s correlation coefficients comparing these TSA-Seq datasets were comparable to those obtained comparing biological replicates (Figure 1H-M).

To better compare SPADs before and after transcriptional inhibition, we performed a percentile normalization of the TSA-Seq data (Figure 1N). Defining SPADs here specifically as genomic regions in the top 5 percentile of SON TSA-Seq scores,^18^ we observed SPADs are largely conserved across all groups: control versus transcription-inhibited cells and log phase versus G1 cells (Figure 1N-P).

We therefore conclude that establishment of NS-association with SPADs does not require ongoing transcription.

### COL1A1 BAC transgenes autonomously target to NS after random integration

Our observations in the previous section demonstrate transcription-independent NS-genome association, as measured by TSA-seq. These results are consistent with an early report of the transcription-independent close association of the *COL1A1* locus with NS.^17^ In contrast, BAC HSPA1 transgenes^32,33^ showed reduced NS-association with transcriptional inhibition. Therefore, using light microscopy we next tested for transcription-independent NS-association for COL1A1 and other gene loci, and if a BAC transgene could recapitulate this transcription-independent NS-gene association.

We used DNA immuno-FISH to validate the NS-association of three genomic loci (GAPDH, COL1A1, COL1A2) (Figure 2A-B) and then to test the sensitivity of their NS-association to transcriptional inhibition (Figure S2). As predicted by TSA-seq (Figure S2A), whereas both GAPDH and COL1A1 showed close NS-association (median distance < 0.23 ± 0.03 µm from NS) in the three human cell types tested (HCT116, hTert-RPE1, hTert-BJ fibroblasts), COL1A2 showed close association only in fibroblasts (Figure 2B). Importantly, the NS-association of all the three gene loci in HCT116 cells was insensitive to transcriptional-inhibition by DRB (Figure S2B-C), consistent with the previously reported transcription-independent NS-association of the endogenous COL1A1 locus.^17^

**Figure 2.**
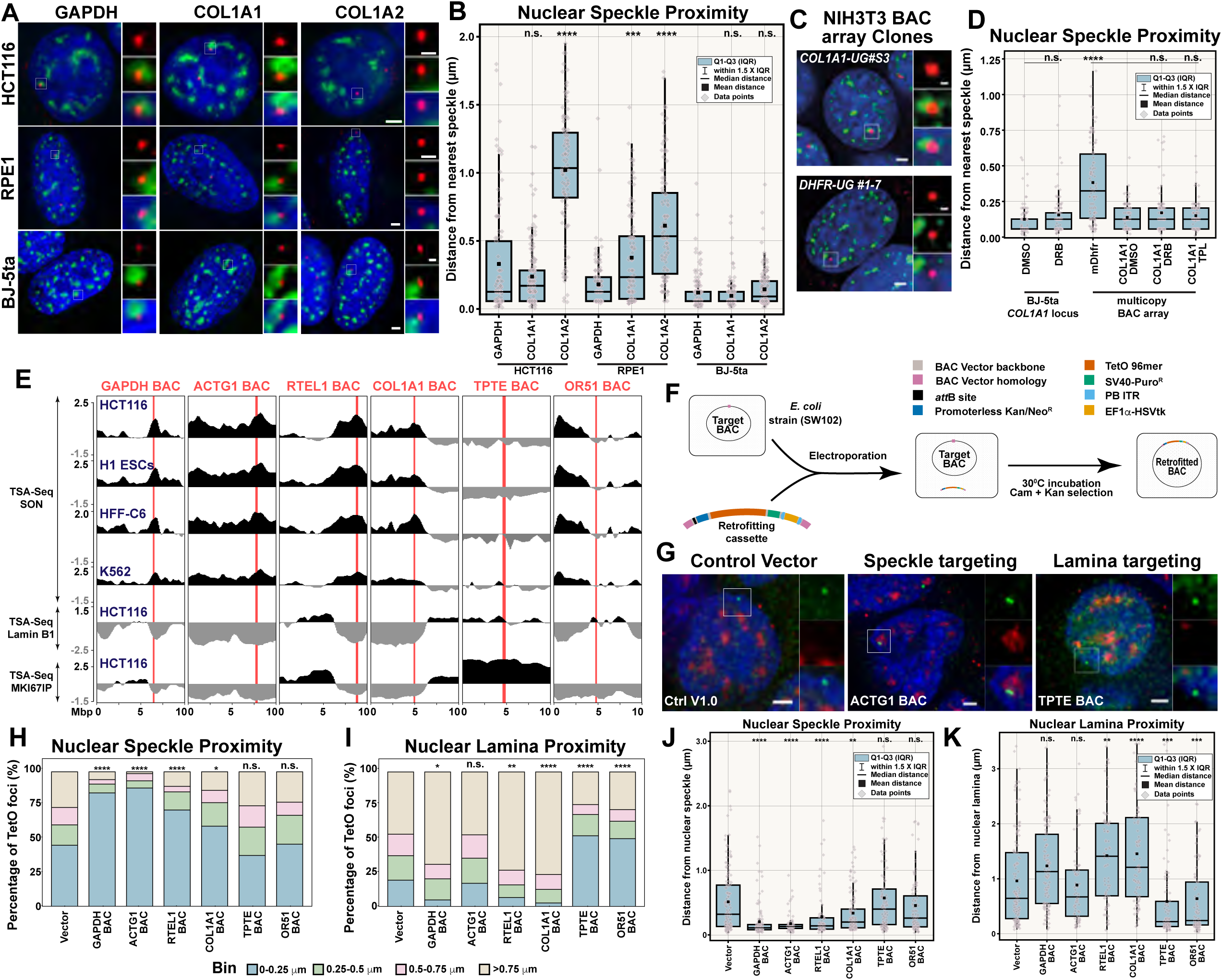
BAC transgenes containing SPAD DNA autonomously target to NS. **(A-B)** Variable NS proximity of endogenous loci. **(A)** Maximum intensity z-projection of 3D DNA FISH (red) for endogenous GAPDH, COL1A1, and COL1A2 genomic loci relative to NS immunostaining (green) and DNA (DAPI, blue) in the indicated cell lines. Scale bars: 2 μm (main panels) and 0.5 μm (insets). **(B)** Box plots showing distances of FISH foci from the nearest NS for loci in (A). Plots display the median (black horizontal line), mean (black square), individual data points (gray diamonds), and 1.5X interquartile range (whiskers). Significance was tested relative to GAPDH loci (n = 104-151). **(C-D)** Transcription-independent NS-targeting of the COL1A1 endogenous locus and BAC transgenes. **C)** Maximum intensity z-projections of 3D DNA FISH (red) for integrated human COL1A1 (Clone UG#S3) and mouse Dhfr (Clone UG#1-7) BACs in NIH 3T3 cells with NS (green) and DAPI (blue). Scale bars as in (A). **(D)** Box plots showing effects of transcriptional inhibition on NS proximity for endogenous COL1A1 locus in BJ-5ta-hTert cells or for multicopy BAC arrays in NIH 3T3 cells. Distances were measured by 3D DNA FISH and compared to DMSO controls (n = 103-106). **(E)** TSA-Seq enrichment profiles for SON (top 4 tracks), lamin B1, and MKI67IP showing proximity of six candidate BACs (highlighted red) to speckles, lamina and nucleoli. **(F)** Schematic of the BAC retrofitting approach. Retrofitting elements (top right) were excised from control plasmid and introduced into the BAC backbone. (G) Maximum intensity z-projections of HCT116 mixed cell populations stably transfected with the indicated control vector or retrofitted BACs. Transgenes are visualized by TetR-EGFP (green), NS (red) and DAPI (blue). Scale bar: 2 μm. **(H-I)** Stacked histograms showing distance distribution of foci to nearest NS **(H)** or nuclear lamina **(I)** for empty vector versus retrofitted BACs. x-axis: construct, y-axis: fraction of foci within each distance bin. Significance was tested by one-tailed z-test for fraction of foci within 0.25 μm (n=74-132). **(J-K)** Box plots of TetO foci distances to NS **(J)** and nuclear lamina **(K)** HCT116 mixed populations containing the indicated constructs. Plots are as in (B). Significance was tested by one-way ANOVA with Tukey’s (HSD) (n = 74-132). n.s., not significant; *P < 0.05; **P < 0.01; ***P < 0.001; ****P < 0.0001.

We next tested whether an integrated multi-copy *COL1A1* BAC transgene array, containing ∼165 kbp of human genomic DNA encompassing and flanking the *PPP1R9B*, *SGCA, COL1A1, TMEM92* genes, would show transcription-independent targeting to NS. We recombineered COL1A1 BAC and derived several stable NIH 3T3 fibroblast cell clones containing integrated copies of the recombineered human COL1A1 BAC transgene using the approach described in our previous study.^34^ We then used immuno-DNA-FISH to measure the NS-association of an ∼1.2 Mbp multicopy COL1A1 BAC transgene array in one of these clones (Clone COL1A1-UG#S3). This multicopy COL1A1 BAC transgene demonstrated strong NS-association in mouse NIH 3T3 fibroblasts (median distance 0.13 ± 0.01 µm from NS), reproducing the NS-association of endogenous COL1A1 locus in human BJ fibroblast (median distance 0.06 ± 0.01 µm from NS) (Figure 2C-D). In contrast, a multi-copy mouse DHFR BAC array of comparable size (∼0.8 Mbp) (Clone DHFR UG#1-7), also containing highly expressed genes, showed a significantly larger distance (p< 0.0001) from NS (median distance 0.33 ± 0.03 µm).

The close NS-association of the COL1A1 BAC transgene array was maintained in the presence of both DRB and TPL (Figure 2D), recapitulating the transcription-independent NS-association of the endogenous COL1A1 locus (Figure S2). We conclude that both the endogenous COL1A1 locus and the COL1A1 BAC transgene show transcription-independent targeting to NS and that therefore BAC transgenes can be leveraged to dissect both transcription-dependent and -independent NS-targeting mechanisms.

### Autonomous Targeting to NS is a general property of BAC transgenes containing SPAD DNA

To test whether autonomous targeting to NS was a general property of SPADs, we next compared the NS versus nuclear lamina targeting of six BAC transgenes containing human DNA inserts of ∼100-200 kbp. As a more rigorous test of autonomous NS-targeting, we decided instead to test the NS-targeting of single-copy BAC transgenes integrated randomly into the genome using piggyBAC (PB) transposition.

We compared the autonomous targeting of single copy insertions of four BACs containing SPAD DNA and two BACs containing DNA from Lamina Associated Domains (LADs). We named the BACs containing human genomic DNA from SPADs after the well characterized genes-GAPDH, ACTG1, RTEL1, and COLA1-contained in each of these human genomic sequences. These “SPAD BACs” each contain human DNA from gene-rich, early-replicating genomic loci (data not shown) located within iLADs and centered at or close to the maxima of broad, Mbp-scale SON TSA-Seq peaks, as measured in multiple cell types (Figure 2E, red lines). As negative controls, we selected two BACs that contain genomic sequences from LADs: one from a mostly constitutive LAD (OR51) and another from a region (TPTE) which also has strong nucleolar association (MKI67IP TSA-Seq) (Figure 2E).

For these experiments, the vector backbone of all the BACs mentioned above was retrofitted by recombineering.^35^ The retrofitting construct adds to the vector backbone a φC31 *attB* site, a neomycin (Neo^R^ ) selectable marker without eukaryotic promoter, a 96mer Tet operator (TetO) array, a SV40 promoter-driven puromycin selectable marker, and an EF1α promoter-driven HSVtK negative-selectable marker flanked by PB inverted terminal repeats (ITRs) (Figure 2F). The retrofitting plasmid (Ctrl V1.0) was included as an internal control to establish the baseline targeting activity of the construct (Figure 2F).

We co-transfected these retrofitted BAC constructs together with a plasmid expressing the PB transposase into HCT116 cells. We established a mixed clonal population of cells selected for BACs integrated via PB transposition rather than nonhomologous end joining (NHEJ) by using an initial puromycin positive selection followed by ganciclovir negative selection. This negative selection eliminates cells containing BAC sequences integrated via NHEJ and thus still containing the HSVtk negative selection marker. However, the HSVtk negative-selectable marker located outside the PB ITRs, does not integrate in host genome in case of PB transposition (Figure 2F, Figure S2D).

To visualize the location of the integrated BAC transgenes, we transduced cells with a lentivirus expressing a Tet repressor-EGFP (TetR-EGFP) fusion protein and fixed cells 5 days later (Figure 2G). All four BAC transgenes containing SPAD DNA showed a statistically significant decreased distance to NS compared to both the LAD-containing BAC transgenes and the empty vector negative controls (Figure 2H&J). Moreover, the NS proximities of these BAC SPAD transgenes were comparable to the NS-proximity of the endogenous COL1A1 and GAPDH genes in HCT116 cells (Figure 2B&J). In contrast, neither of the LAD BAC transgenes showed a statistically significant change in NS-proximity compared to the vector-only negative control. Instead, both LAD BAC transgenes showed closer proximity to the nuclear periphery than the empty vector negative control (Figure 2G, I&K). Moreover, three of the four BAC SPAD transgenes showed a decreased proximity to the nuclear periphery relative to the vector only negative control (Figure 2I&K), suggesting that these BAC SPAD transgenes are primarily moving away from the nuclear periphery and towards NS, known to be located towards the nuclear interior.^6,36^

In summary, these results demonstrate that autonomous NS-targeting is a common property of ∼100-200 kbp fragments of DNA derived from SPAD regions.

### Functionally redundant NS-targeting cis elements exist within the COL1A1 BAC

Having established the autonomous NS-targeting activity as a common property of SPAD-containing BAC transgenes, we next used this targeting activity as an assay to identify smaller cis-elements. We focused on the COL1A1 BAC primarily because the COL1A1 locus was earlier shown to have transcription-independent NS-targeting,^17^ which based on our TSA-Seq results appears to be the dominant mode of NS-targeting genome-wide. Additionally, it had the lowest gene density of the four SPAD BAC transgenes we tested, which we hypothesized might correlate with a lower density of NS-targeting cis-elements, thereby facilitating our dissection of the NS-targeting activity of the BAC.

Because the endogenous COL1A1 locus showed its closest proximity to NS in fibroblasts of the three cell lines tested, we decided to perform our assay in fibroblasts for maximum sensitivity. We used mouse NIH 3T3 fibroblasts because of their high transfection efficiency of large BAC transgenes,^34^ and also because we would be able to distinguish human COL1A1 transgene sequences from the endogenous mouse COL1A1 sequences.

We continued our use of PB transposition for this dissection of the COL1A1 BAC. Smaller fragments of the COL1A1 BAC in a vector backbone containing the PB ITRs, the TetO 96mer array, and a Zeocin selectable marker were created through an iterative BAC recombineering approach.^35^ These fragments, ranging in size from 11-159 kbp, spanned most of the 165 kbp COL1A1 genomic insert in the original COL1A1 BAC (Figure 3A). Using the same PB transposition approach described in the previous section, we created mixed clonal cell populations containing integrated BAC transgenes for each of the constructs containing the smaller COL1A1 fragments shown in Figure 3A.

**Figure 3.**
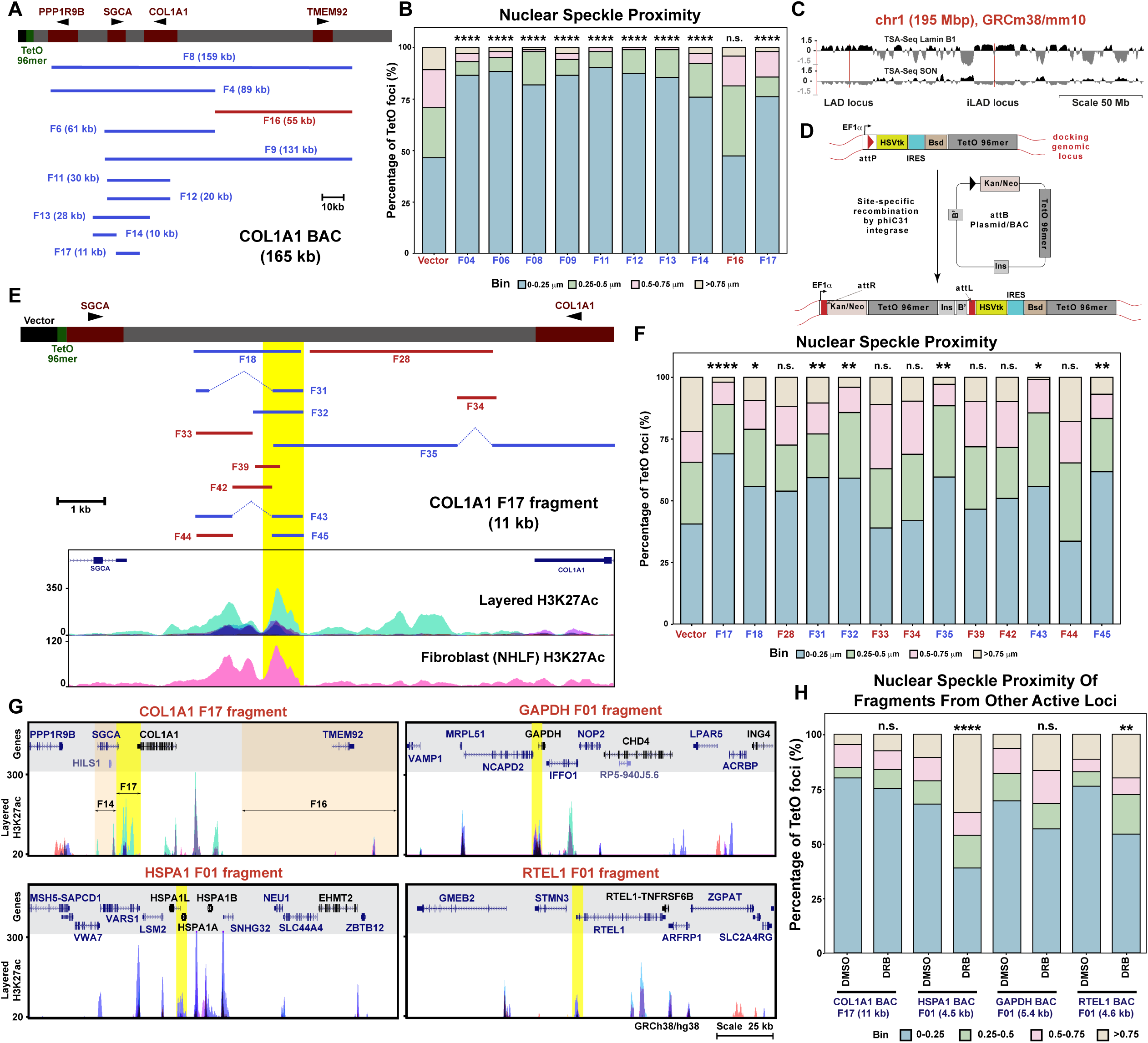
Identifying minimal NS-targeting sequences within the COL1A1 BAC and other SPADs. **(A)** Schematic of the COL1A1 BAC showing genes (red rectangles with arrowheads indicating the direction of transcription), a TetO-96mer array (green), and vector backbone (black). Solid lines indicate BAC fragments with sizes in parenthesis; Fragments with statistically significant (blue) and non-significant (red) NS-targeting are shown. **(B)** Stacked histogram showing distances of TetO foci to the nearest NS for PB-mobilized empty vector or indicated BAC fragments in NIH 3T3 cells. x-axis: construct; y-axis: percentage of foci within each bin. Significance was tested by one-tailed z-test for the fraction of foci within 0.25 μm (n=97-105). **(C)** Genome browser view of mouse chromosome 1 showing Lamin B1 and SON TSA-Seq profiles with selected regions (red) used for CRISPR-Cas9 docking line generation. **(D)** Schematic of the approach used for site-specific integration of NS-targeting sequences in the COL1A1 BAC. Only assay-relevant elements are shown. **(E)** Schematic showing the positions of tested BAC fragments within the larger COL1A1 F17 fragment. Blue fragments showed statistically significant NS-targeting activity; red fragments showed non-significant differences from control. Yellow highlight marks overlap with layered H3K27ac peaks across seven cell types including normal human lung fibroblast (NHLF). **(F)** Stacked histogram showing distances of TetO foci to the nearest NS for the empty vector or indicated BAC fragments integrated at the iLAD locus in NIH 3T3 cells. Significance tested by one-tailed z-test for foci within 0.25 μm (n=93-104). **(G)** Genome browser view of four genomic loci overlayed with H3K27ac profiles. Highlighted peaks (yellow) were subcloned and tested for NS-targeting activity. Stacked histogram showing distances of TetO foci to the nearest NS for the empty vector or indicated BAC fragments integrated at the iLAD locus in NIH 3T3 cells following DMSO or DRB treatment. Significance tested by one-tailed z-test for foci within 0.25 μm (n= 60-105). n.s., not significant; *P < 0.05; **P < 0.01; ***P < 0.001; ****P < 0.0001.

Initial dissection of COL1A1 BAC from its 3’-end demonstrated comparable NS-targeting of a smaller BAC transgene (F4) to the NS-targeting of a near full-length F8 BAC fragment (Figure 3A-B). We then demonstrated similar NS-targeting of additional smaller BAC fragments, F6 and F9, made by the deletion of a common 5’ region of fragments F4 and F8. Additional deletions from the 5’-end or 3’-ends led us to two non-overlapping 11 kbp fragments, F14 and F17, with comparable NS-targeting to that of the near full-length F8 fragment (Figure 3B). On the contrary, the 55 kbp F16 fragment covering much of the 3’ half of the COL1A1 BAC did not show any NS-targeting (Figure 3B).

Because F14 and F17 do not overlap but individually show NS-targeting comparable to that of the near full-length COL1A1 BAC, we conclude that there are at least two functionally redundant NS-targeting cis elements within the COL1A1 BAC.

### An ∼600 bp region within the F17 COL1A1 fragment targets to NS

We decided to further dissect one of the two 11 kb NS-targeting fragments, F17, containing the intergenic region between the SGCA and COL1A1 genes.

In the earlier experiments we scored NS-targeting activity for different COL1A1 BAC fragments in mixed clonal populations, assuming random PB transposition at different genomic locations. Integration of 10s-100s of kbp of BAC transgene can shield the influence of neighboring chromatin,^28,34,37,38^ but smaller DNA fragments might be more prone to chromosome position effect. Therefore, through a site-specific recombination approach, we analyzed the NS-targeting activity of progressively smaller fragments integrated at the same genomic location.

We adopted φC31 integrase mediated site-specific recombination at specific genomic loci in mouse NIH 3T3 fibroblasts (Figure 3C-D, Figure S3A). The φC31 integrase enables precise, unidirectional integration of incoming construct through recombination between the bacteria-derived attB and phage-derived attP sequences.^39,40^ This approach of integrating constructs through attP/attB-based site-specific integration is more suited than traditional CRISPR-based approach to integrate constructs larger than 5 kb.^41^ We first generated docking lines containing the attP site, flanked by an EF1α promoter regulated HSVtK counter-selectable marker and blasticidin (Bsd) positive selectable marker. We then used site-specific integration via the φC31 integrase to integrate sequences from attB-containing plasmid or BAC constructs (Figure 3D). Successful integration places the Kan/Neo selectable marker under the control of the EF1α promoter, enabling its expression, while concurrently displacing the HSVtK and Bsd markers from the promoter, thereby abolishing their expression (Figure 3D). The inactivation of HSVtK expression with successful integration allows the use of ganciclovir counterselection to increase the fraction of colonies corresponding to true site-specific recombinants.

We used an engineered NIH 3T3 cell clone (#3C4) containing a docking site inserted into a small iLAD region (Reg 03, Figure S3A). We chose this iLAD location to insert the docking site because TSA-seq showed this chromosome region to be at intermediate distances away from both NS and the nuclear lamina (Figure S3). This docking site was created by CRISPR/Cas9-mediated targeted insertion of the attP-HSVtK-IRES-BSD^R^ cassette containing a 96mer TetO array. The full-length F17 fragment, and smaller fragments derived from the F17 fragment, were cloned into the Ctrl V2.0 vector used for site-specific recombination. We then performed attP-attB recombination to derive NIH 3T3 cell clones containing F17 or smaller fragments in this docking site. Individual clones containing the integrated COL1A1 fragments were genotyped to verify site-specific recombination.

Using this attP-attB targeting approach, we measured F17 NS-targeting activity (Figure 3F) that was comparable to the F17 NS-targeting activity, previously observed using PB random integration (Figure 3B). Analysis of the NS-targeting of the smaller F17 fragments revealed multiple fragments with statistically significant NS-targeting activity. All these NS-targeting fragments shared a smaller common region (Figure 3E, yellow highlighted region, Figure 3F). The smallest of these fragments, F45, is ∼600 bp and had NS-targeting activity only marginally lower but not statistically different than the full-length 11 kb F17 fragment. Moreover, the NS-targeting activity of the F45 fragment was comparable to other larger fragments which overlapped with F45 (Figure 3F).

Thus, we were able to reconstitute essentially the full NS-targeting capability of the 11 kb F17 fragment with a sequence as small as ∼600 bp. This targeting activity was not as strong as that of the full-length COL1A1 BAC but was comparable to the NS-targeting activity of the 11 kb F14 fragment, located 5’ of the F17 fragment (Figure 3A-B). We speculate that multiple, partially functionally redundant NS-targeting fragments contribute to the NS-targeting activity of the full-length COL1A1 BAC.

### NS-targeting activity is a common property of other several kbp cis elements with elevated H3K27ac

From our analysis of the 165 kbp COL1A1 locus (Figure 3A-B), we identified two 11 kbp fragments (F14, F17) with NS-targeting activity and one 55 kbp fragment (F16) without NS-targeting activity. Interestingly, both 11-kbp fragments overlap with several kbp-wide high-amplitude peaks of H3K27ac present in multiple cell lines, including human fibroblasts. However, no such high-amplitude peak was observed within the 55 kbp fragment which did not show NS-targeting activity (Figure 3G). Additionally, the ∼600 bp F45 NS-targeting DNA fragment contained within the F17 fragment encompasses a subpeak of H3K27ac (Figure 3E). A similar H3K27ac peak in CUT&Tag profiles is also evident at the F45 fragment syntenic genomic region of immortalized mouse embryonic fibroblast (iMEF) cells (data not shown).^42^

Large regions of elevated H3K27ac (∼10-60 kbp) are a hallmark of super-enhancers, previously shown to be enriched near SON TSA-seq peaks.^18,43^ Given the overlap of NS-targeting fragments with peaks of H3K27ac, we next asked whether NS-targeting activity might be a common property of other several kbp cis elements with elevated H3K27ac. We hypothesized that larger regions of elevated H3K27ac with properties of super-enhancers, likely to contain several enhancers, would contain at least one NS-targeting cis-element within them, consistent with the previously described enrichment of super-enhancers in top deciles of SON TSA-seq.^18^

To test this possibility, we cloned three ∼5-6 kb-size DNA fragments overlapping with such H3K27ac peaks from three different BAC transgenes shown here (GAPDH, RTEL1, Figure 2) and elsewhere (HSPA1)^28^ with NS-targeting activity: GAPDH F01 (5.8 kb), RTEL1 F01 (4.8 kb), and HSPA1 F01 (5 kb) (Figure 3G). These fragments were integrated into the iLAD attP docking locus in NIH 3T3 cells and tested, together with the 11 kbp F17 COL1A1 fragment, for their NS-targeting activity both in the absence and presence of the transcriptional inhibitor, DRB (Figure 3H).

All three of these fragments showed significant NS-targeting activity relative to the Ctrl V2.0 vector-only negative control (Figure 3H). We previously discussed the transcription-independent NS-targeting of the COL1A1 BAC versus the reported transcription-dependent NS-targeting of the HSPA1 BAC. Interestingly, for these ∼5-6 kbp fragments, we also observed both transcription-independent (GAPDH F01 and COL1A1 F17) and transcription-dependent (RTEL1 F01, and HSPA1 F01) NS-targeting (Figure 3H).

We conclude that NS-targeting is a common property of several kbp cis elements with elevated H3K27ac modification. Moreover, consistent with previous observations, these smaller cis elements also display two different NS-targeting activities: one that is transcription-dependent and another that is transcription-independent.

### A subset of transcription factor motifs provides redundant NS-targeting activity to the F45 fragment

We next sought to identify the specific cis regulatory elements (CREs) responsible for the NS-targeting activity of the F45 ∼600 bp DNA fragment. We began by focusing on known transcription factor (TF) binding motifs within this fragment, as predicted by the JASPAR 2022 Transcription Factor Binding Site Database (Minimum Score set at 500, *p* value < 10^-5^),^44^ hosted on the UCSC genome browser.^45^ Five sites of predicted transcription factor binding motifs were predicted; as a group, these included potential binding motifs for 8 different transcription factors (Figure 4A). Four of these transcription factors (TFAP2A, TFAP2B, TFAP2C, and MXI1) were predicted to bind to overlapping motifs contained within the most 5’ site. In the middle of the F45 fragment, the next site corresponds to overlapping predicted binding sites for ZBTB18 and TWIST1, followed by a site containing a single binding site for ZNF384. Towards the 3’ end of the F45 fragment is a site with a single binding site for ZNF460, followed by a cluster of overlapping ZNF384 binding sites at the 3’ end.

**Figure 4.**
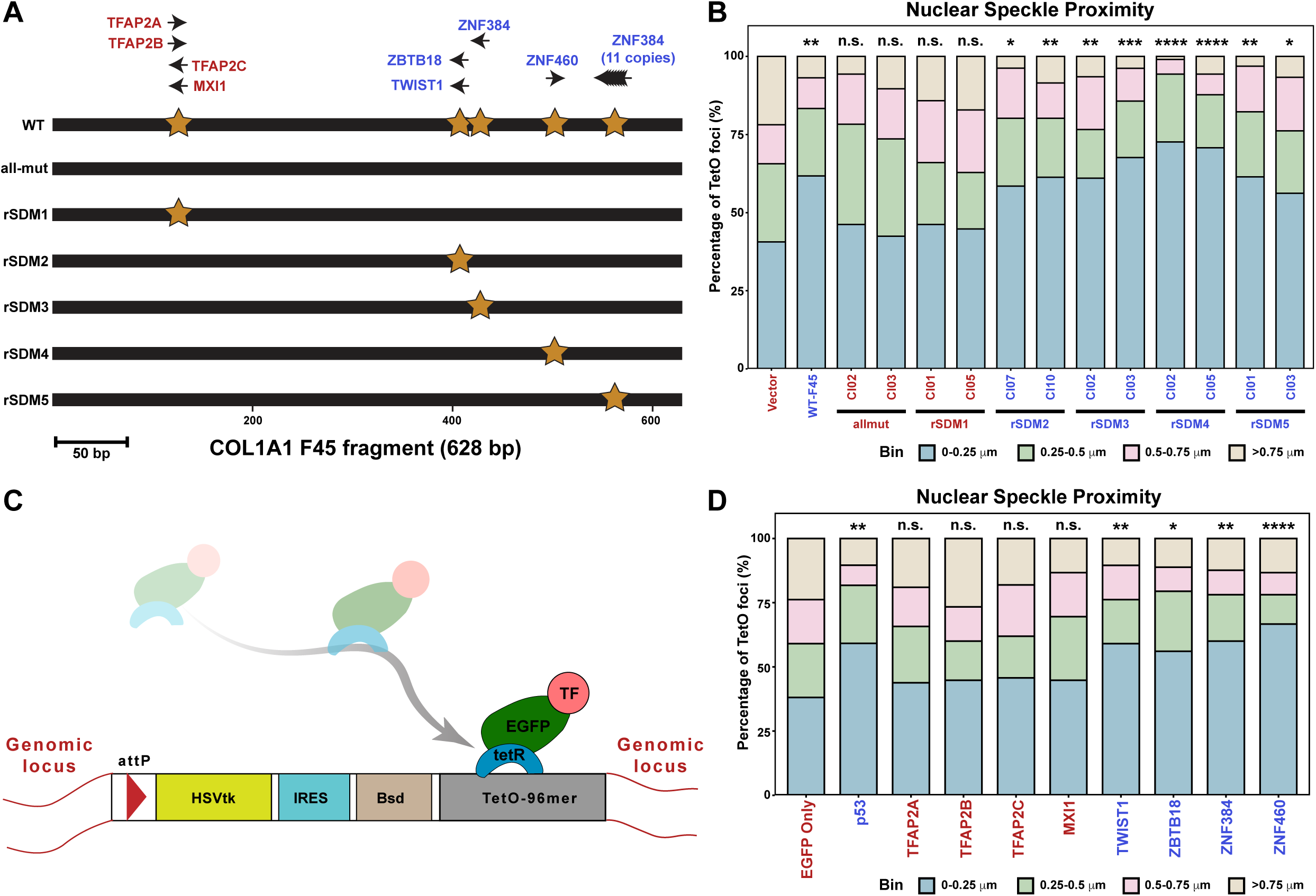
Identification of transcription factor binding motifs conferring NS-targeting activity within the ∼600 bp COL1A1 BAC F45 fragment. **(A)** Schematic of the 600 bp F45 fragment with predicted transcription factor (TF) binding sites. Predicted TF motifs are indicated by brown stars, with arrows showing motif orientation. Mutant versions of F45 used for dissecting NS-targeting are also shown. Top: Blue-labeled TFs are predicted to bind to motifs that conferred NS-targeting; red-labeled TFs are predicted to bind to motifs that did not confer statistically significant NS-targeting. **(B)** Stacked histogram showing distances of TetO foci to the nearest NS for empty vector or the indicated constructs integrated at the iLAD locus in NIH 3T3 cells. x-axis: construct; y-axis: percentage of foci within each distance bin. **(C)** Schematic of the TetR-EGFP-TF protein recruitment assay. Fusion proteins are recruited to the locus via TetO-TetR interaction. **(D)** Stacked histogram showing distance distribution of TetO foci to the nearest NS for the indicated fusion proteins recruited to the iLAD locus in NIH 3T3 cells. Significance tested by one-tailed z-test for foci within 0.25 μm (n= 105-115). n.s., not significant; *P < 0.05; **P < 0.01; ***P < 0.001; ****P < 0.0001.

We tested if these 5 identified sites for TF binding were together necessary and sufficient for NS-targeting activity of the F45 fragment by measuring the NS-targeting of a mutated (“all-mut”) F45 fragment. This all-mut F45 fragment was created by mutating all predicted TF binding motifs contained within the original F45 fragment. To minimize possible changes to nucleosome positioning and/or the relative positioning or rotational positioning of binding sites for unidentified trans factors, we used nucleotide substitutions rather than nucleotide deletions within the core TF binding motifs (Figure S4). After integrating this all-mut F45 fragment into the iLAD docking site, we observed that the measured NS-association was not statistically different from the negative empty vector control (Ctrl V2.0), as confirmed in two independently derived cell clones with this all-mut F45 integration (Figure 4A-B).

We, therefore, concluded that one or more of these TF binding sites are necessary and sufficient for the NS-targeting activity of the F45 fragment. We next asked which if any of these TF binding sites were capable of independently conferring NS-targeting activity.

A new set of mutated F45 fragments was constructed by restoring each single TF binding site separately to the all-mut F45 fragment, creating 5 mutated F45 fragments, each containing just one wild-type TF binding site (Figure 4A, rSDM1 through rSDM5 fragments). After site-specific integration we isolated two NIH 3T3 cell clones for each of these mutated F45 fragments, corresponding to independent integrations into the docking site, and measured NS-targeting of these fragments in both of these cell clones.

Restoring the overlapping binding motifs for the TFAP2A/B/C and MXI1 TFs in the 5’ end of the all-mut F45 fragment (rSDM1 fragment) failed to rescue the NS-targeting activity observed for the wild-type F45 fragment; the NS-targeting of the rSDM1 fragment was not significantly different from the vector negative control (Figure 4B). Therefore, we conclude that this site, containing possible binding sites for TFAP2A/B/C and MXI1, is not capable of independently targeting to NS.

In contrast, restoration of the remaining TF binding motif sites in DNA fragments rSDM2-5 rescued the NS-targeting activity to levels comparable to the wild-type F45 fragment in each of the two independently isolated cell clones tested for each fragment (Figure 4B). We therefore conclude that each of the four different TF binding motif sites in the F45 fragment is individually sufficient to target to NS with comparable efficiency as the original F45 fragment. These binding motifs potentially recruit ZBTB18, TWIST1, ZNF384 and ZNF460 TFs (Figure 4A). Notably, the rSDM3 fragment containing one ZNF384 binding motif showed similar NS-targeting activity to the rSDM5 fragment containing 11 potential ZNF384 binding sites.

Overall, these results demonstrate a functional redundancy of NS-targeting activities contained within this single ∼600 bp DNA F45 fragment that is distributed over four out of the five distinct TF factor binding sites predicted for this fragment.

### A subset of the TFs predicted to bind to the F45 fragment show NS-targeting activity in a tethering assay

In the preceding Section, we identified a subset of TF binding motifs that individually could rescue the NS-targeting activity of the ∼600 bp F45 fragment from the COL1A1 BAC. In some cases, more than one TF was predicted to bind to the same binding motif or overlapping binding motifs.

To test whether the NS-targeting activity could be assigned to individual TFs, we next developed a tethering approach to separately assay the NS-targeting activity of each of these TFs (Figure 4C). The attP/attB site-specific recombination approach that we used introduced a total of 192 TetO repeats-96 introduced during creation of the attP docking line and another 96 with the attB insertion plasmid (Figure 3D). The presence of the second 96mer TetO repeat was meant to enable a visual validation of cell clones with true site-specific integration of the attB plasmid. Therefore, we used fusions of TetR-EGFP with each TF for our tethering assay and only TetR-EGFP as a negative control for this assay. A prior study had reported NS-association of p53 transcription factor through its proline-rich domain.^24^ Therefore, we used TetR-EGFP fused with the full-length p53 as a positive control.

One technical difficulty with this approach was that many of the TetR-EGFP-TF fusion proteins showed non-diffuse, localized nucleoplasmic distribution, even in the absence of any TetO repeat. This non-diffuse background binding made identification of the tagged docking site difficult to distinguish from all other sites of localized binding. This problem did not exist for the negative control TetR-EGFP, which showed diffuse background binding. We overcame this problem by instead using DNA FISH to detect the TetO repeats when TetR-EGFP TF fusion proteins were tested.

We again used the attP docking site inserted into the small iLAD (Figure S3A). Using this approach, we show a statistically significant increased NS-targeting after expression of the TetR-EGFP-p53 fusion protein relative to the TetR-EGFP negative control (Figure 4D), consistent with the previously reported NS-targeting activity of p53.^24^ Additionally, we show NS-targeting activity for the full-length TWIST1, ZBTB18, ZNF384, and ZNF460 TFs (Figure 4D). In contrast, tethering of TFs TFAP2A/B/C and MXI1 showed no statistically significant difference in NS-targeting as compared to the TetR-EGFP negative control (Figure 4D).

Thus, each TF predicted to bind to the 4 TF binding sites with NS-targeting activity within the F45 fragment conferred NS-targeting activity comparable to p53 in this tethering assay. In contrast, none of the four TFs predicted to bind to the rSDM1 TF binding site showed NS-targeting activity in this tethering assay.

### Targeting to NS is a common property of acidic activation domains, including those found in NS-targeting TFs with binding motifs in the COL1A1 F45 fragment

Previously, the observation that mutations in a proline-rich domain of p53 decreased NS-targeting of the p53 TF led to the suggestion that multiple TFs likely target to NS through a common proline-rich domain.^24,46^ However, there was no clear correlation between the presence or absence of proline-rich domains with the presence or absence of NS-targeting among the 8 TFs identified in our dissection of the F45 COL1A1 DNA fragment. Instead, we found that the 4 out of 8 TFs with NS-targeting activity (TWIST1, ZBTB18, ZNF384, ZNF460) had been previously annotated as containing acidic activation domains (AAD), thus being defined as acidic transcription factors.

AADs contain intrinsically disordered activation domains, enriched in acidic residues flanking clusters of hydrophobic amino acids, especially aromatic and leucine residues.^47–50^ Interestingly, our laboratory previously demonstrated that tethering the strong VP16 AAD to a heterochromatic, amplified chromosome region led to large-scale chromatin decondensation^49^ and induced long-range, directional movement of a plasmid transgene from the nuclear periphery towards the nuclear interior.^51^ This large-scale chromatin decondensation activity was common property of several AADs, including a synthetic peptide ‘DELQPASIDP’ (DELQ peptide),^52,53^ but not for glutamine-rich or proline-rich non-acidic transcriptional activation domains (NAADs). Subsequent studies demonstrated that the recruitment of VP16, p65 AAD, DELQ peptide, and VP64-a tetramer of VP16, can also lead to relocalization of other endogenous chromosome loci from the nuclear periphery to the interior. ^54,55^ Further, while studying chromatin decondensation of the heterochromatic, amplified chromosome region by VP16 recruitment, we noticed clustering of NS around the periphery of the amplified region.^49^

We therefore decided to ask whether AADs have NS-targeting activity as a common property. We first started with previously well characterized AADs (Figure S5) for which we had previously measured large-scale chromatin decondensation (VP16, p65 AAD1 and AAD2, DELQ peptide, p53) ^52,53^ and also, in some cases, chromosome repositioning activity (VP16, DELQ peptide, p65 AAD2) using lac-repressor tethering assays.^51,52,54^ For comparison, we also measured the NS-targeting activity of other previously tested glutamine-rich (SP1, OCT1, OCT2) and proline-rich (AP-2A, also known as TFAP2A) transcription activation domains from nonacidic transcription factors, collectively referred here as NAADs (Figure S5). Additionally, we also measured the NS-targeting activity of the N-terminal domain of CTCF, which previously had been reported to have large-scale chromatin decondensation activity using the same tethering assay as used for VP16;^56^ we note that CTCF also recently has been suggested to strengthen the ground-state NS association of NS-proximal genes.^46^

To visualize the location of the TetO repeat, we transiently expressed TetR-EGFP fused to each of these protein domains for five days after lentivirus delivery in the NIH 3T3 cell clone containing the iLAD attP docking site with a TetO 96-mer repeat. Strikingly, both the VP16 AAD and DELQ peptide conferred strong NS-targeting activity relative to the TetR-EGFP control (VP16, 70.7% <0.25 μm from NS, mean 0.23 ± 0.02 μm, *p =* 0.0013; DELQ peptide, 71.7% <0.25 μm from NS, mean 0.23 ± 0.02 μm, *p*=0.0010). The other AADs (p65 AAD1, p65 AAD2, p53 AAD) also conferred statistically significant NS-targeting activity relative to the TetR-EGFP negative control (Figure 5B, left histograms). In contrast, NS-targeting was not significantly different from the TetR-EGFP negative control either for the tested NAADs or for the N-terminal domain of CTCF (Figure 5B, right histograms).

**Figure 5.**
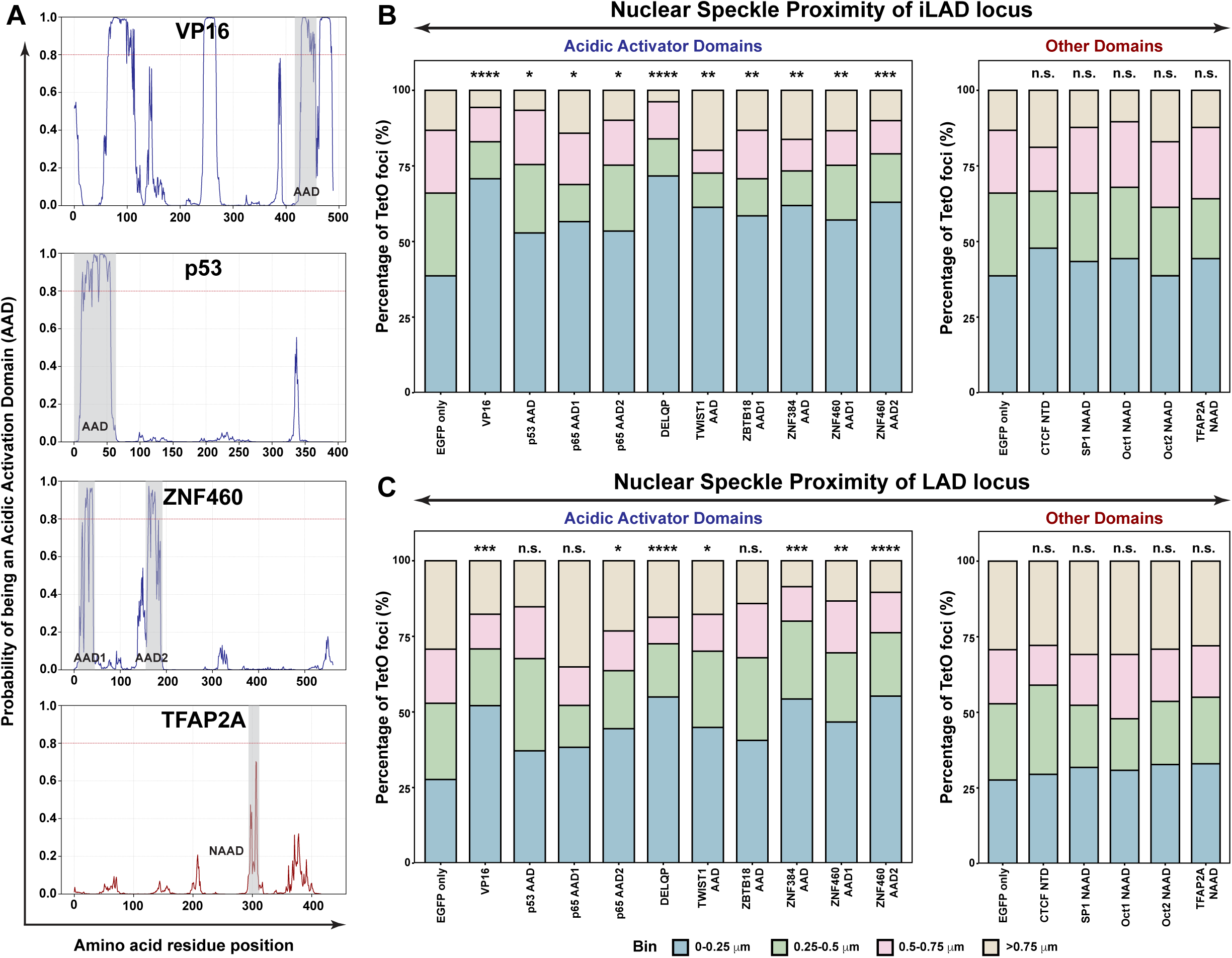
**Acidic Activator Domains (AADs) show NS-targeting activity that is chromatin context dependent**. (A) Representative plots of predicted acidic transcription activation domains (AADs) in the indicated transcription factors. Regions tested in the TetO-TetR recruitment assay are shaded in gray. The red horizontal lines indicate the AAD prediction threshold. (B-C) AADs show NS-targeting activity, but not other activation domains (NAADs) or the CTCF N-terminal domain. Stacked histogram showing distances of TetO-96mer signals to the nearest NS for the TetR-EGFP fusion proteins recruited to the NIH 3T3 iLAD locus (B) or LAD docking sites (C). x-axis: Protein domains fused to TetR-EGFP; y-axis: percentage of foci within each distance bin. Significance was tested by one-tailed z test of proportions within 0.25 μm (n= 61-123). n.s., not significant; *P < 0.05; **P < 0.01; ***P < 0.001; ****P < 0.0001.

Having established a common NS-targeting activity for AADs as a group, we proceeded to ask whether the AADs within the acidic transcription factors identified as trans factors for the NS-targeting activity of the F45 COL1A1 fragment (ZNF384, ZNF460, ZBTB18, TWIST1) also shared this NS-targeting activity and, if so, how the NS-targeting activity of these AADs compared to the activity of the full-length acidic transcription factors. We selected AAD-like peptides from the four NS-targeting acidic TFs through a deep-learning model ADpred, trained to identify AAD positive sequences, using its default settings (minimum score ≥0.8 over ≥10-15 continuous residues). For comparison, we applied the same AAD prediction model as well to the 4 other TFs predicted to bind to the F45 COL1A1 fragment but without measurable NS-targeting activities (TFAP2A/B/C and MXI1), as well as to all the TFs containing the other peptides already tested for their NS-targeting activities, as described earlier in this Section.

As predicted by this model, transcription factors ZBTB18, ZNF384, and ZNF460 have peptide sequences with scores predicting them to be AADs. These scores were comparable to the predicted ADpred scores for the previously studied NS-targeting AADs (VP16 AAD, p65 AAD1 and AAD2, p53 AAD) (Figure 5A, Figure S5A, grey highlights). TWIST1 had a peptide sequence with a moderate ADpred score, but not quite high enough to be classified by this model as an AAD. TFAP2A and TFAP2B had a similar peptide sequence with low predicted AAD scores, noticeably lower than the scores of the TWIST1 peptide sequence. All other TFs examined had no peptide sequences with moderate to high scores in this AAD-prediction model. (Figure S5A). Based on this model’s predictions, we chose peptide sequences (Figure S5A grey highlights, Figure S5B) to test NS-targeting activity using the same TetO/TetR tethering assay.

In line with the predictive model scoring of potential AADs, all predicted AADs (ZBTB18 AAD, ZNF384 AAD, ZNF460 AAD1 & AAD2) as well as the TWIST1 peptide sequence which showed moderate to high values of the AAD-predictive score showed NS-targeting activity comparable to the previously tested AADs from VP16, p65, and P53 (Figure 5B). In contrast, all other tested peptide sequences with lower AAD-predicted scores, including the peptide sequence from TFAP2A, showed no significant NS-targeting. This was consistent with the previously demonstrated lack of NS-targeting activities of the entire TF sequences from which these peptides were derived (Figure 4).

In conclusion, our results demonstrate first that NS-targeting activity is a common property of AADs; no AAD tested failed to show NS-targeting activity. Second, of all TFs identified as having binding motifs within the NS-targeting F45 COL1A1 DNA fragment, only the TFs with NS-targeting activity contained AAD-like sequences capable of conferring NS-targeting activity. Moreover, this AAD NS-targeting activity was comparable to that observed by the full-length parent TFs.

### NS-targeting activity of AADs is chromatin context-dependent

In our original experiment showing repositioning of a plasmid transgene from the nuclear periphery to the nuclear interior after lac repressor-VP16 tethering,^51^ we looked for but failed to observe a gain in NS association (unpublished observation). For this reason, we wondered if there might be a chromatin context dependence for our observed AAD targeting to NS. In all the previous NS-targeting assays, we used a docking site located in an iLAD.

To address this question, we compared the NS-targeting of TetO repeats inserted into a large LAD (Reg 01, Figure S3) versus the iLAD docking site after tethering of AADs and other transcription factor domains described earlier (Figure 5B). We observed significantly reduced NS-targeting for the tested AADs when they were instead tethered to the same TetO 96mer located within the LAD as compared to when they were tethered to the iLAD (Figures 5B-C). The AAD NS-targeting remained statistically significant but was reduced in magnitude for most of the tested AADs (VP16, DELQ peptide, p65 AAD2, TWIST1, ZBTB18, ZNF384, ZNF460 AAD1 & AAD2). For the p53 AAD and p65 AAD1 AADs, NS-targeting activity within the LAD was higher but not statistically different from that observed with the TetR-EGFP negative control.

In conclusion, while NS-targeting is a common property of AADs, the magnitude of their NS-targeting activity can be influenced by the chromatin-context. This observation predicts that there will not be a direct correlation between the binding of acidic transcription factors across the genome and NS proximity.

### NS-targeting by AADs is transcription-dependent

Given that for different DNA fragments (Figure 3G-H) we observed two types of NS-targeting activities, transcription-dependent and transcription-independent, we explored whether NS-targeting conferred by tethered AADs was sensitive to transcriptional inhibition. We compared the NS-association of our two strongest targeting fusion proteins, containing VP16 or DELQ peptide, tethered both at the iLAD and LAD docking sites in the presence or absence of TPL (Figure 6A-B). More specifically, we first compared the NS-association of these AADs after TPL treatment relative to their NS-association with the solvent control (DMSO) to determine if there was a statistically significant decrease in NS-association. Second, we compared the NS-association after TPL treatment for each of the AADs relative to the TetR-EGFP negative control treated similarly with TPL. These two comparisons together were expected to reveal if transcription-inhibition causes reduced NS-targeting by the AADs or if it completely eliminates NS-targeting to the levels that are not significantly different from the similarly treated TetR-EGFP control.

**Figure 6.**
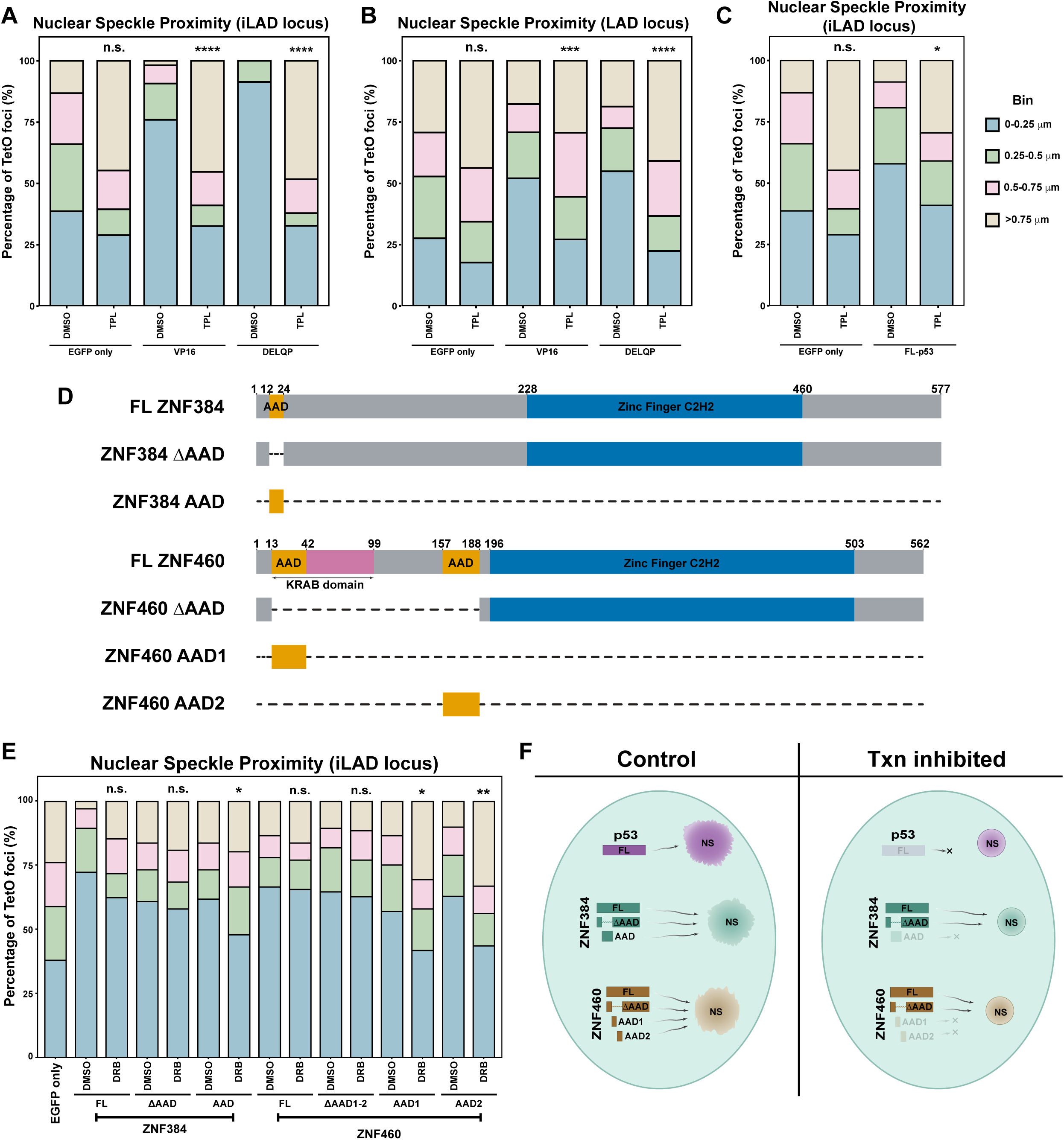
Some acidic transcription factors show both transcription-dependent NS-targeting via their acidic activatior domains (AADs) and transcription-independent NS-targeting through additional domains. **(A-B)** Stacked histograms showing distances of TetO-96mer signals to the nearest NS for TetR-EGFP fusion AADs recruited to the NIH 3T3 iLAD locus (A) or LAD locus (B) following DMSO or triptolide (TPL) treatment (n= 54-123). (C) Stacked histogram showing distances of TetO-96mer signals to the nearest NS for TetR-EGFP alone or TetR-EGFP-p53 (full-length) recruited to the iLAD locus after DMSO or triptolide (TPL) treatment. (n= 57-106). (D) Schematic of individual TF domains and deletion constructs. Dotted line indicates deleted residues; double-headed arrows mark overlapping domains in ZNF460. Numbers denote amino acid boundaries. (E) Stacked histogram showing distances of TetO-96mer signals to the nearest NS for TetR-EGFP fusion proteins recruited at the NIH 3T3 iLAD locus after DMSO or DRB treatment (n= 96-105). (F) Summary schematic of NS-targeting by TetR-EGFP fusion constructs at the iLAD locus after DMSO, DRB, or TPL treatment. Solid-colored constructs indicate significant NS-association; lighter shades indicate loss of NS-targeting under the conditions tested. For all the stacked histograms (panels A-C, andE): x-axis: construct; y-axis: percentage of foci within each distance bin. Significance was tested by one-tailed z test of proportions within 0.25 μm, n.s., not significant; *P < 0.05; **P < 0.01; ***P < 0.001; ****P < 0.0001.

For all of these experiments, we used the TetR-EGFP signals to visualize distances from NS. We observed that after TPL treatment, the TetR-EGFP negative control at both tested loci show a small but statistically insignificant decrease in NS-association (Figure 6A-B). We suspect that the presence of the selectable marker used for creating the docking lines is providing a weak and transcription-dependent NS-targeting activity.

Our results show that after TPL transcriptional inhibition, the VP16 AAD and AAD-like DELQ peptide show a decreased NS-targeting activity at the iLAD docking site, which was indistinguishable from the TetR-EGFP negative control (Figure 6A). Similar results were observed as well for the LAD docking site (Figure 6B). These results demonstrate that, in general, the NS-targeting by AADs is transcription-dependent.

### Some acidic transcription factors contain additional domain(s) conferring transcription-independent NS-targeting activity

The transcription-dependent NS-targeting of AADs described in the previous Section created a paradox. Our TSA-seq experiment showed that most genomic regions established their NS-association independent of transcription (Figure 1). We went on to demonstrate how COL1A1 BAC transgenes recapitulated this transcription-independent NS-targeting (Figures 2 and 3). Our dissection of the COL1A1 F45 DNA fragment led us to correlate acidic TFs with NS-targeting. However, we then demonstrated that NS-targeting is a common property of AADs but that this AAD NS-targeting activity may be transcription-dependent.

To address this paradox, we tested if other domains exist within acidic TFs that provide transcription-independent NS-targeting.

While this work was in progress, another study associated NS-targeting activity with a proline-rich domain (PRD) within p53.^24^ To test whether such an additional domain might confer transcription-independent NS-targeting by p53, we repeated the TPL inhibition experiment using a TetR-EGFP fusion protein with the full-length p53. In this case we used DNA FISH for TetO spot localization. After transcriptional inhibition by TPL, we observed a statistically significant reduction of NS-association compared to NS-targeting in presence of DMSO solvent control. Further, this loss of NS-targeting by full-length p53 was comparable to the NS-targeting observed for TetR-EGFP control treated with TPL (Figure 6C). Therefore, we conclude that the full-length p53 and any NS-targeting domains within it show transcription-dependent NS-targeting.

We next repeated similar analyses for the ZNF384 and ZNF460 TFs that showed the most statistically significant NS-targeting activity among the 4 TFs with NS-targeting activity predicted to bind to the F45 COL1A1 fragment. Moreover, we compared the NS-targeting activities of the full-length versions of these two TFs to the corresponding deletions without their AADs. Because ZNF460 has two AADs, we made a larger deletion that eliminates both AADs (Figure 6D).

Relative to the full length ZNF384, the ΔAAD ZNF384 deletion showed no statistical difference in NS-targeting activity with or without DRB inhibition, whereas the ZNF384 AAD showed no statistical difference in NS-targeting activity without DRB but a decrease in NS-targeting activity with DRB treatment (Figure 6E). Similar results were observed for ZNF460. After deletion of both AADs, both the full-length ZNF460 and the double ΔAAD ZNF460 showed NS-targeting activities that were not significantly different with or without DRB. However, each ZNF460 AAD separately showed NS-targeting activity comparable to the full length ZNF460, but a decrease in NS-targeting activity with DRB treatment.

In conclusion, both ZNF384 and ZNF460 show a transcription independent NS-targeting activity. This is independent of their AADs, but must depend on additional, yet to be identified protein domains that confer a NS-targeting activity similar to that of the two full-length TFs and the AADs contained within these TFs.

## DISCUSSION

The proximity of highly active genes to nuclear speckles (NS) has led to a transcription-driven model of NS association,^57^ whereby the observed transcription-dependent concentration of splicing factors and other RNA-binding proteins (RBPs) over the transcripts produced by highly active gene loci in both telophase and interphase nuclei then leads to a nucleation of NS containing many of these same RBPs.^3,25,58,59^ However, conclusions from SPRITE, TSA-Seq, multiplexed FISH, and super-resolution imaging reveal that only a subset of highly active genes are stably NS-associated,^18,19,21,22^ with a large fraction of highly active chromosomal loci not closely associated with NS.^36^ Conversely, the COL1A1 locus remains NS-associated even after transcriptional inhibition,^17^ indicating that factors beyond transcription must play critical roles in genome positioning relative to NS.

Therefore, here we set out to identify determinants of the highly reproducible positioning of Speckle Associated Domains (SPADs) adjacent to NS using a top-down, unbiased approach. At the genome-wide scale, using TSA-seq, we first demonstrated that active transcription is neither required to maintain NS-chromatin associations in interphase nuclei nor to establish new contacts during *de novo* speckle formation in post-mitotic nuclei. In contrast to previous suggestions and models,^3,25,57^ these observations suggest that the close NS-association of large, 100s-1000s kbp SPADs with high gene density is not simply a downstream consequence of high levels of gene expression. This presents a fundamental question: if transcription is dispensable, what other mechanisms may determine and/or redundantly reproduce basal NS-targeting?

We addressed this question by random integration of ∼150-200 kbp DNA sequences selected from the centers of SPADs, followed by dissection of a ∼165 kb COL1A1 BAC, a locus representing a prominent subpeak within a several Mbp sized SPAD. Our data indicate that rather than large contiguous genomic stretches, NS-targeting is primarily determined by multiple, functionally redundant, smaller intrinsic cis-regulatory DNA elements (CREs) present within tested SPADs. Two smaller, ∼11 kb NS-targeting fragments identified within the COL1A1 BAC corresponded to genomic sites enriched in H3K27ac. Integration of several 5-6 kb DNA fragments overlapping such H3K27ac peaks established that NS-targeting was a common property of several kbp sequences containing broad peaks of H3K27ac enrichment, a mark of super-enhancers.^18,43^

Our results further demonstrate that functional redundancy in NS-targeting extends to the level of 100s of base-pairs. For instance, an ∼600 bp COL1A1 BAC fragment contained multiple binding motifs for acidic TFs, each independently sufficient to induce NS-targeting comparable to the wild-type fragment. Even these acidic TFs could carry two distinct NS-targeting activities localized to different protein regions: a transcription-dependent NS-targeting activity localized to their AADs and a transcription-independent NS-targeting activity localized elsewhere in a subset of acidic TFs.

This high degree of redundancy similarly accounts for the demonstrated fine-tuning of NS association by the additional binding of individual acidic TFs such as p53 and HIF2α.^24,46^ Physiologically, signal-induced activation of p53 or HIF2α triggers their binding to a limited number of sites, adding their NS-targeting activities to a larger number of constitutively-bound NS-targeting TFs, and consequently leading to a modest repositioning of target genes closer to NS and a concomitant increase in their expression. Even small positional shifts relative to NS can yield multi-fold increase in gene expression through gene expression amplification with nuclear speckle contact.^20,24,46,60^ We found, however, that the NS-targeting activity of acidic TFs was strongly chromatin-context dependent, with reduced NS-targeting activity within LAD compared to iLAD regions. This may help explain why genes moving closer to NS after HSF1, p53, or HIF2α induction are frequently located in genomic regions already prepositioned near NS.^20,24,46^

The complete spectrum of protein domains with NS-targeting activities remains unidentified. A previous study attributed such activity for p53 exclusively to its proline-rich domain, as deletion of these domains reduced targeting, while AAD mutations disrupting transcriptional activation had no effect.^24^ This study did not test the p53 AAD specifically for its targeting activity because of its assumption that this NS-targeting activity would be dependent on its transcriptional activation activity. However, here we showed that AAD-targeting activity can be separate from its transcriptional activity, as seen for the AAD-like DELQ peptide that strongly targets to NS but does not show transcriptional activation. Other proteins with similar proline-rich domains have been proposed to mediate NS-targeting, including HIF2α.^46^ Of the 8 TFs predicted to bind the 600 bp COL1A1 NS-targeting fragment, the four acidic TFs all show NS-targeting activity in our assays, but only one (ZNF384) contained a predicted proline-rich speckle targeting motif.^46^ Conversely, of the four nonacidic TFs, which did not show NS-targeting activity in our assays, three contain a predicted proline-rich speckle targeting motif (TFAP2A/B/C).^46^

Several additional proteins have been suggested to have NS-targeting activity, including the CTCF-related boundary-element and cohesin binding protein MAZ, CTCF, and cohesin, which was suggested to confer NS-targeting via a proline-rich speckle targeting motif in RAD21.^46,61^ By defining NS hubs through inter-chromosomal (trans) Hi-C interactions, it was found that the degree of NS-association showed a strong correlation with trans Hi-C contacts mapping to constitutive super-enhancer sequences rather than H3K27ac levels or super-enhancer activity.^61^ This is consistent with our experimental observations showing NS-targeting activity of 4/4 super-enhancer like DNA fragments (5-6 kbp in size) with broad peaks of elevated H3K27ac, as well as our observation, though, that not all regions with elevated H3K27ac target to NS. Prior examination of the NS-proximal sequences showed high enrichment for MAZ whose knockdown led to decreased NS interactions for strongly NS-associated loci, as validated by microscopy, but increased interactions for weakly NS-associated loci.^61^

The magnitude of the NS-targeting activities of CTCF and cohesin and how they relate to the NS-targeting activities of acidic TFs remains unclear for several reasons.^46^. First, by microscopy, the shifts in NS-positioning induced by CTCF or cohesin, observed using high statistical power, were small and occurred only for genomic regions already positioned close to NS. This contrasts with the ability of AADs to confer long-range shifts in chromosome positioning from NS-distal sites near the nuclear periphery to NS-proximal positioning towards the nuclear interior (data not shown).^51,54^ Second, shifts in NS-targeting were primarily inferred indirectly using molecular proximity assays (CUT&RUN or CUT&Tag) for SON, likely measuring local accumulation of SON over the DNA that appear as sharp, several hundred bp wide peaks centered over CTCF sites.^27,46^ We did not observe any NS-targeting activity for N-terminal domain of CTCF, which was previously shown to drive large-scale chromatin decondensation similar to the VP16 AAD^52^ and to recruit cohesin.^62^

Summarily, our findings establish a multifactorial functionally redundant framework of intrinsic CREs and recruited proteins that cooperatively position specific genomic loci around NS. Such a “Velcro-like” multifactorial NS-targeting model (Figure 7) explains both the robustness of NS–genome interactions across cell types and their plasticity in facilitating rapid gene regulation.

**Figure 7.**
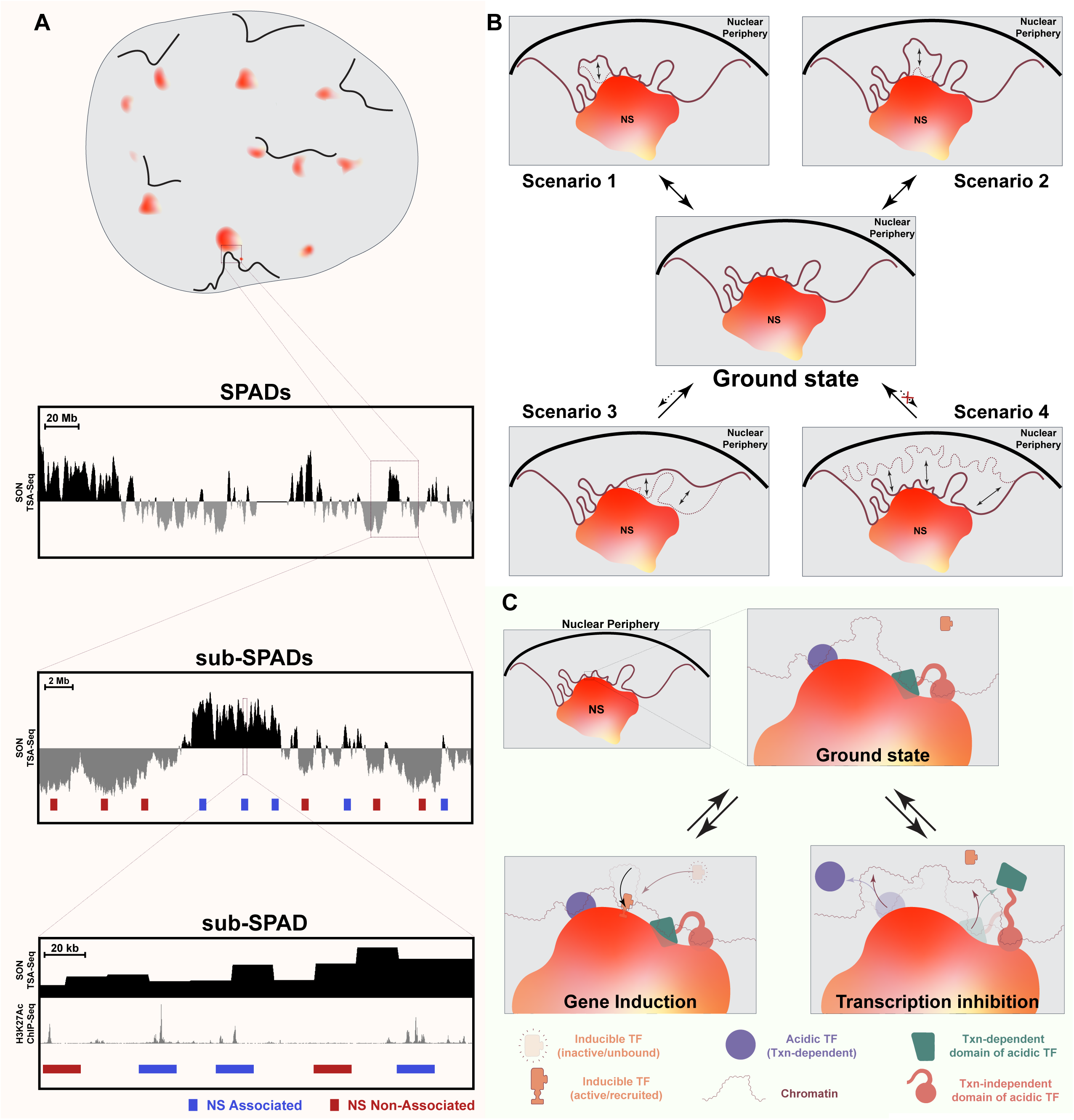
Model of cooperative and redundant mechanisms anchoring genomic loci at the NS periphery. Schematic illustrating how distinct, partially redundant factors act together to position genomic loci at the NS periphery. Both transcription-dependent and transcription-independent transcription factors contribute to stable anchoring at the NS periphery. In this model, acidic activator domains promote transcription-dependent targeting, while additional TF domains and other unidentified elements provide transcription-independent recruitment. The combined action of these pathways ensures robust and cooperative NS association across diverse genomic loci.

### Future directions

Moving forward, several key questions remain. We observe a strong correlation between NS-targeting activity and the presence of H3K27ac peaks, yet only a subset of hyperacetylated sites conferred NS-association. It remains to be tested whether selective removal or pharmacological disruption of H3K27ac peaks alters NS-targeting. More broadly, general mechanisms of transcription-dependent and transcription-independent NS-targeting remain unclear. Recruitment of AADs can generate long-range chromosome movements,^51^ raising the possibility that their NS-targeting is linked to this chromosome movement. If true, this would imply that a distinct mechanism provides stable NS association or “sticking”, potentially via cooperative recruitment of other NS-targeting proteins and/or protein domains-including for example the proline-rich NS-targeting domain described elsewhere,^24,46^ or expressed nascent RNAs at the flanking genes. This may at least partially explain the chromatin-context dependence, such as the reduced NS-targeting activity when AADs were tethered to a TetO repeat inserted within a gene-poor LAD. Chromatin context of NS-targeting was also demonstrated previously in a more physiological context by the dependence of NS-association of the beta-globin locus on its genomic surroundings.^58^ Finally, we speculate that additional protein domains outside of the AADs in acidic TFs or entirely different NS-targeting proteins may also impart transcription-independent NS-targeting.

Another open question concerns implications of NS proximity for gene expression in health and disease. NS-targeting by p53 and HIF2α has been studied in detail.^24,46^ Interestingly, the NS-targeting ZNF384 TF identified in our COL1A1 BAC dissection contains both an N-terminal AAD as well as a C2H2 domain that scaffolds DNA damage repair complexes.^63^ ZNF384 regulates expression of matrix metalloproteinases (MMPs),^64^ and cyclin D1;^65^ overexpression of both has been implicated in cancer progression and invasion. Additionally, ZNF384 is over-expressed in acute leukemias and is specifically enriched at promoters, enhancers and TAD boundaries.^66^ Importantly, over 19 fusion oncoproteins involving ZNF384 with partners such as EP300, TAF15, CREBBP, or ARID1B are found in B-cell acute lymphoblastic leukemia (B-ALL).^67^ It is important to investigate whether these fusions misdirect epigenetic regulators to NS-associated loci or reposition genomic loci bound by ZNF384 fusion partners to NS, thereby causing aberrant transcription that drives leukemogenesis.

Finally, our recent work identifying perispeckle networks as additional niches for active transcription,^68^ suggests a possible continuum of NS targeting. We propose that weaker NS-targeting DNA elements may position loci at low frequency to NS but at higher frequency to these perispeckle networks away from NS. Indeed, fewer super-enhancers map near weaker, “type II” SON TSA-seq peaks, compared to the greater number of super-enhancers observed near SPADs.^18^ Future studies will be needed to dissect these gradations of targeting, including contributions from expressed RNAs and cooperative factors. Addressing these questions may refine distinctions between basal NS anchoring versus dynamic repositioning relative to both NS and these perispeckle networks while clarifying how each contributes to the regulation of gene expression.

## STAR METHODS

Detailed methods are provided in the online version of this paper and include the following:

- **KEY RESOURCES TABLE**
- **EXPERIMENTAL MODEL DETAILS**

**a. CELL LINES**

**METHOD DETAILS**

a. **PLASMID DETAILS**
b. **TRANSIENT TRANSFECTION, DRUG TREATMENT**
c. **GENERATION OF LENTIVIRAL PARTICLES**
d. **GENERATION OF STABLE AND KI LINES**
e. **IMMUNOFLUORESCENCE**
f. **FLUORESCENCE IN SITU HYBRIDIZATION (FISH)**
g. **TSA-SEQ**

**QUANTIFICATION AND STATISTICAL ANALYSIS**

**a. IMAGE ANALYSIS**

## KEY RESOURCES TABLE

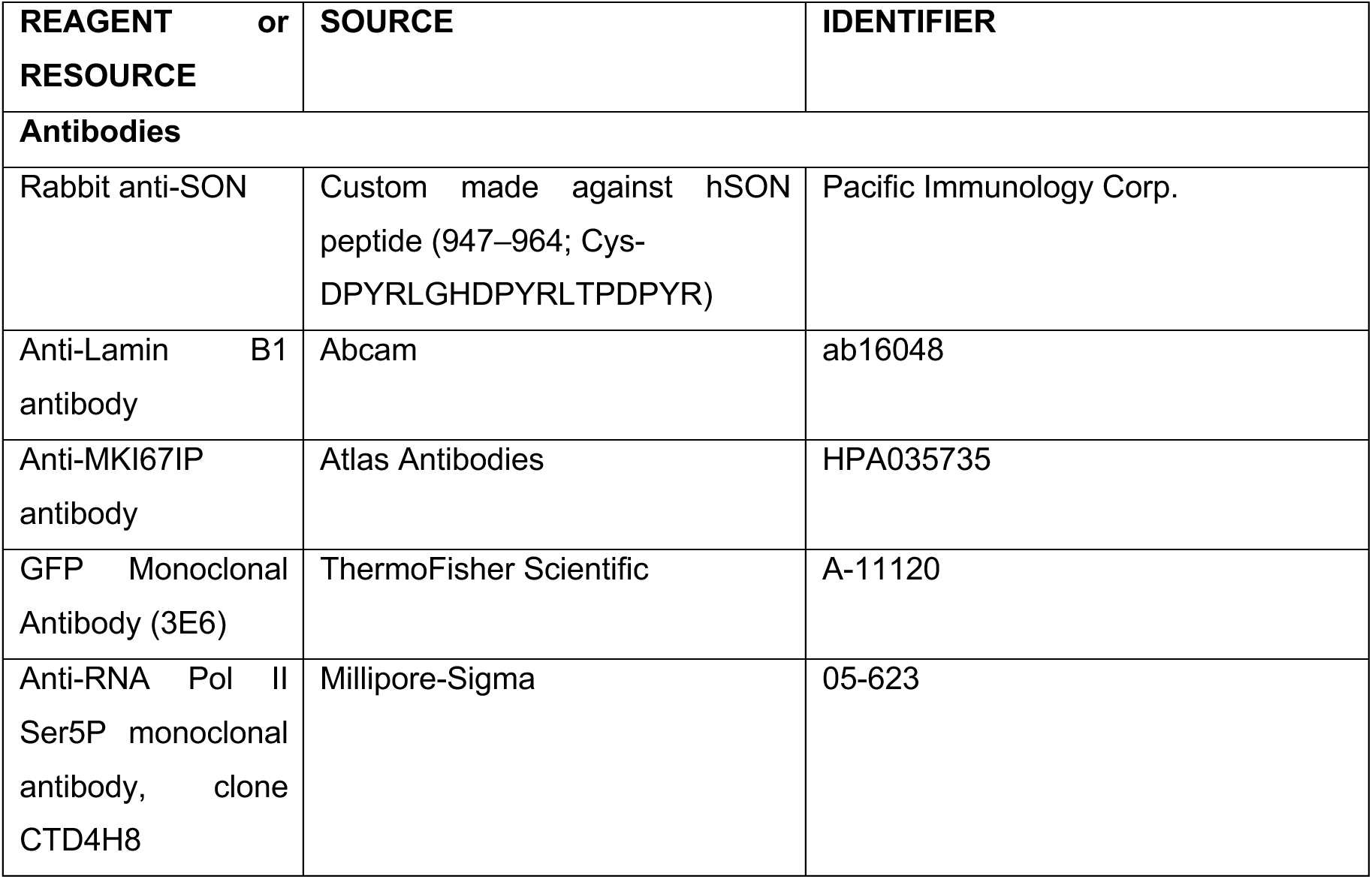
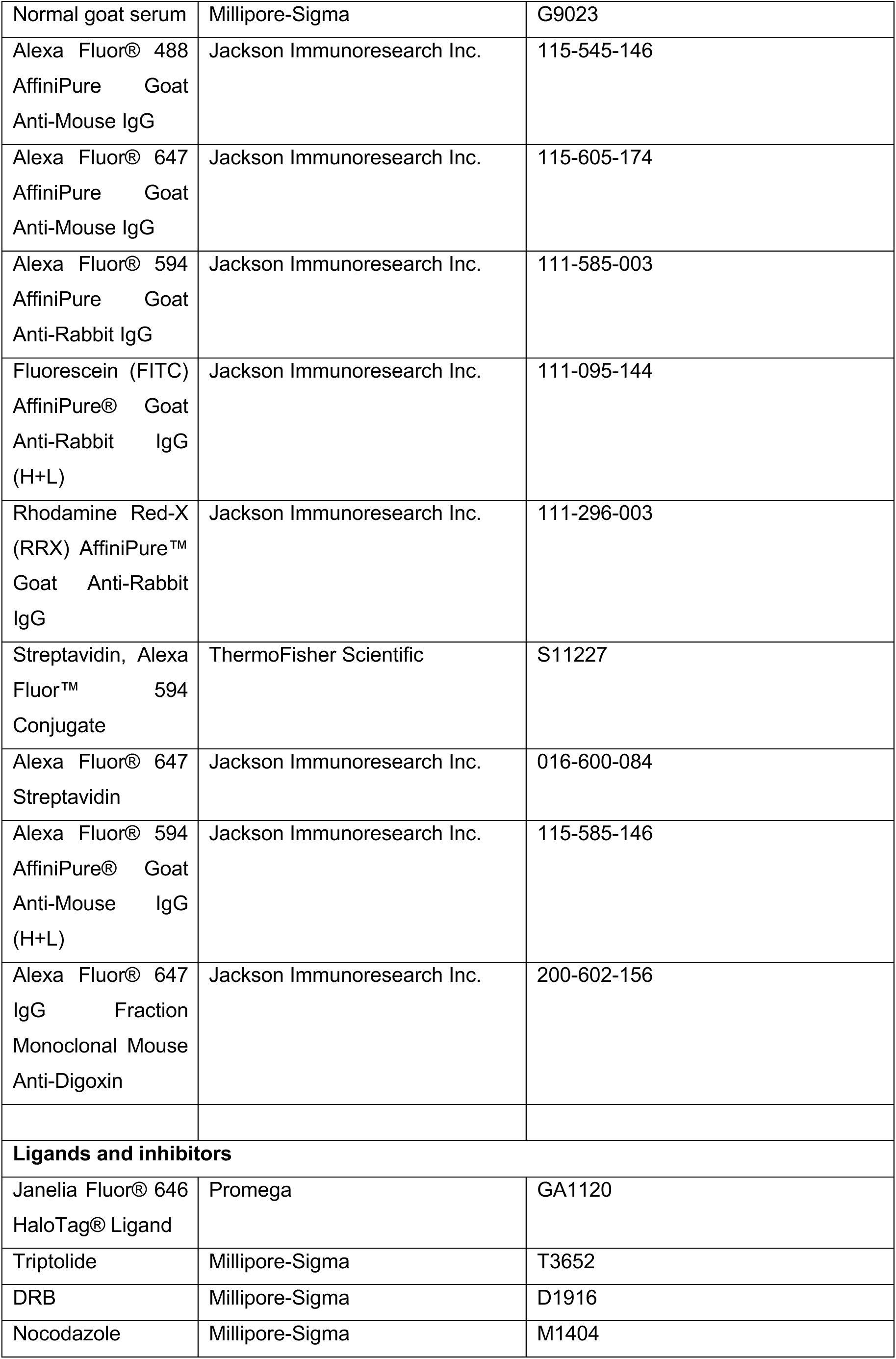
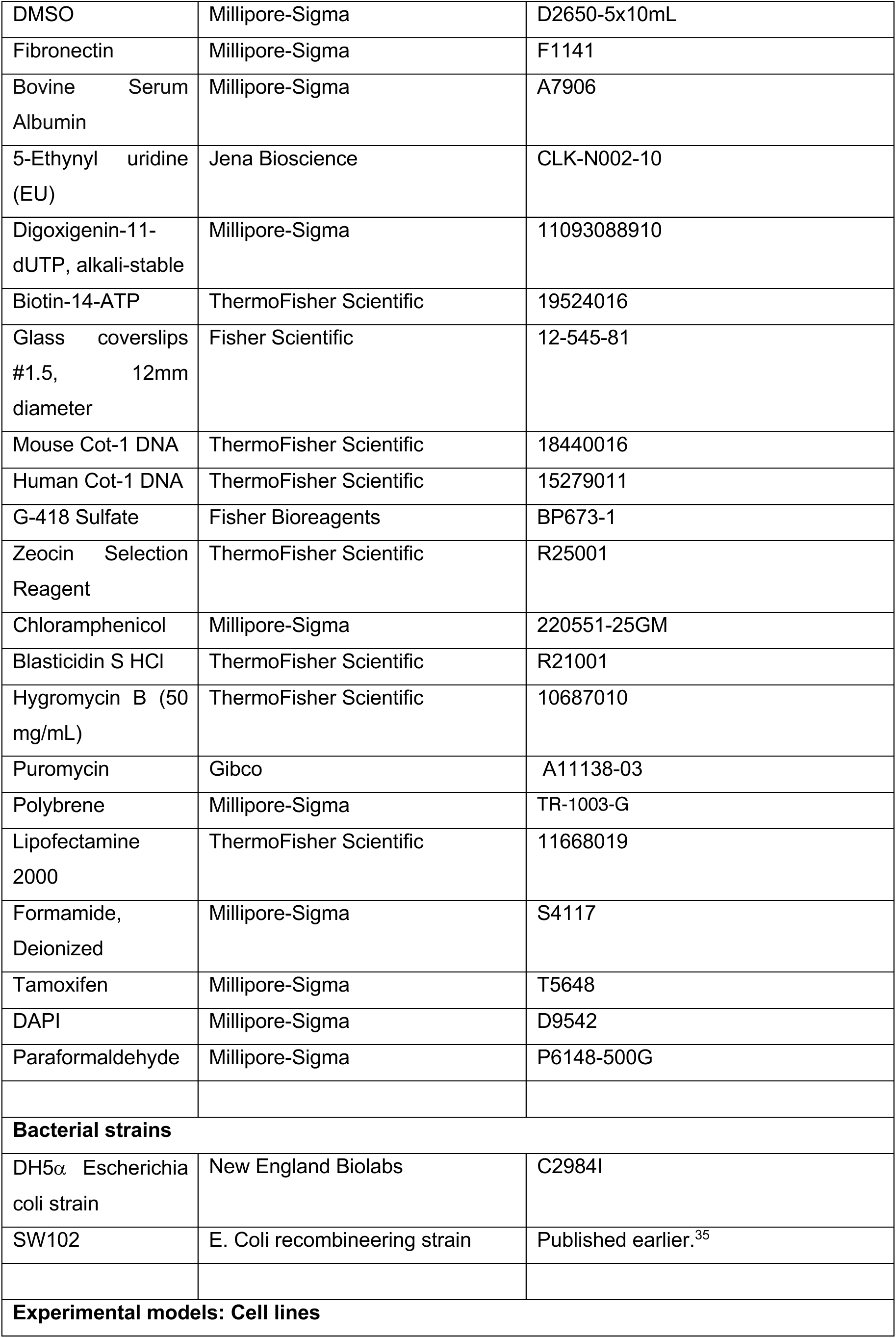
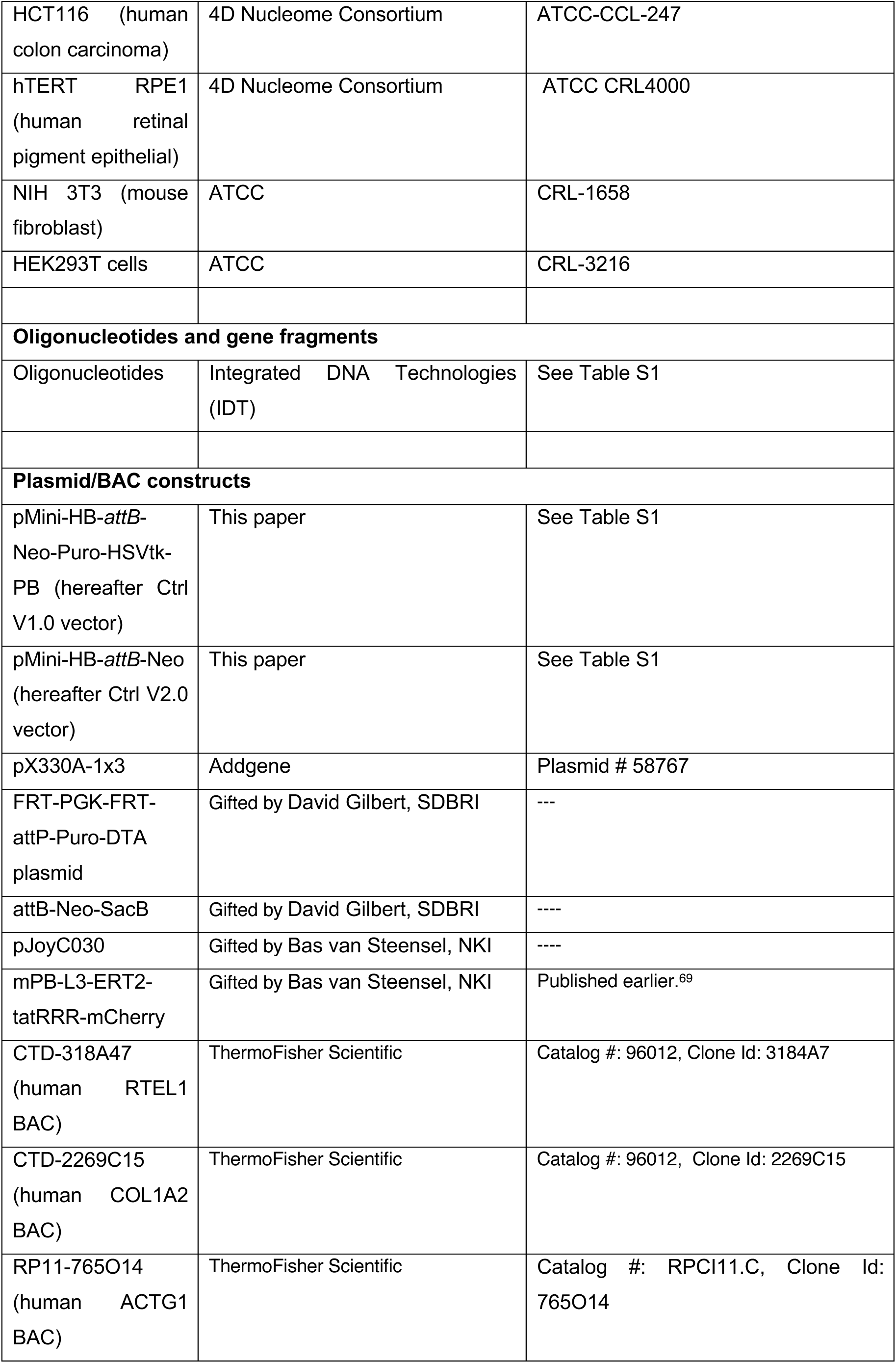
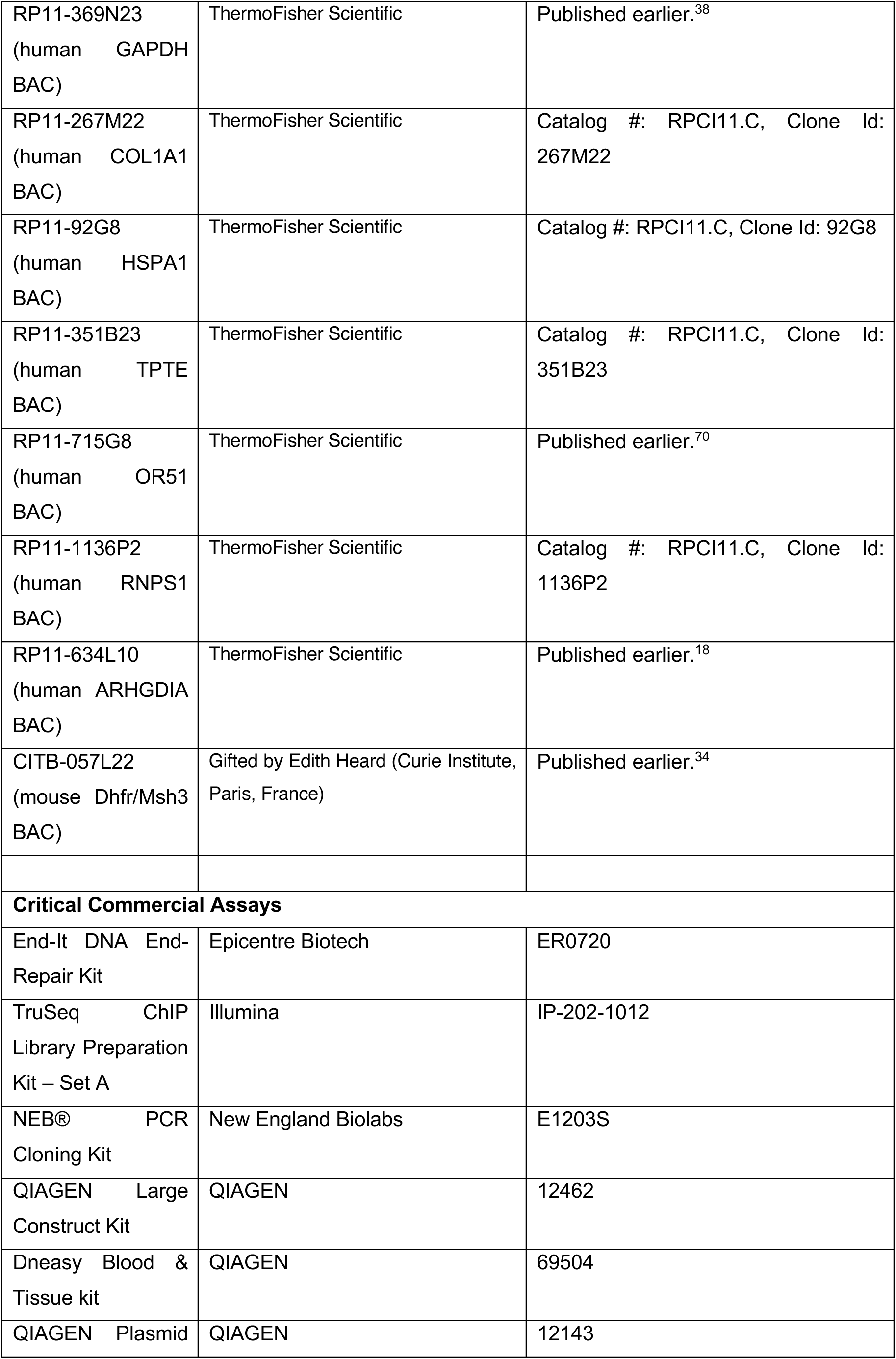
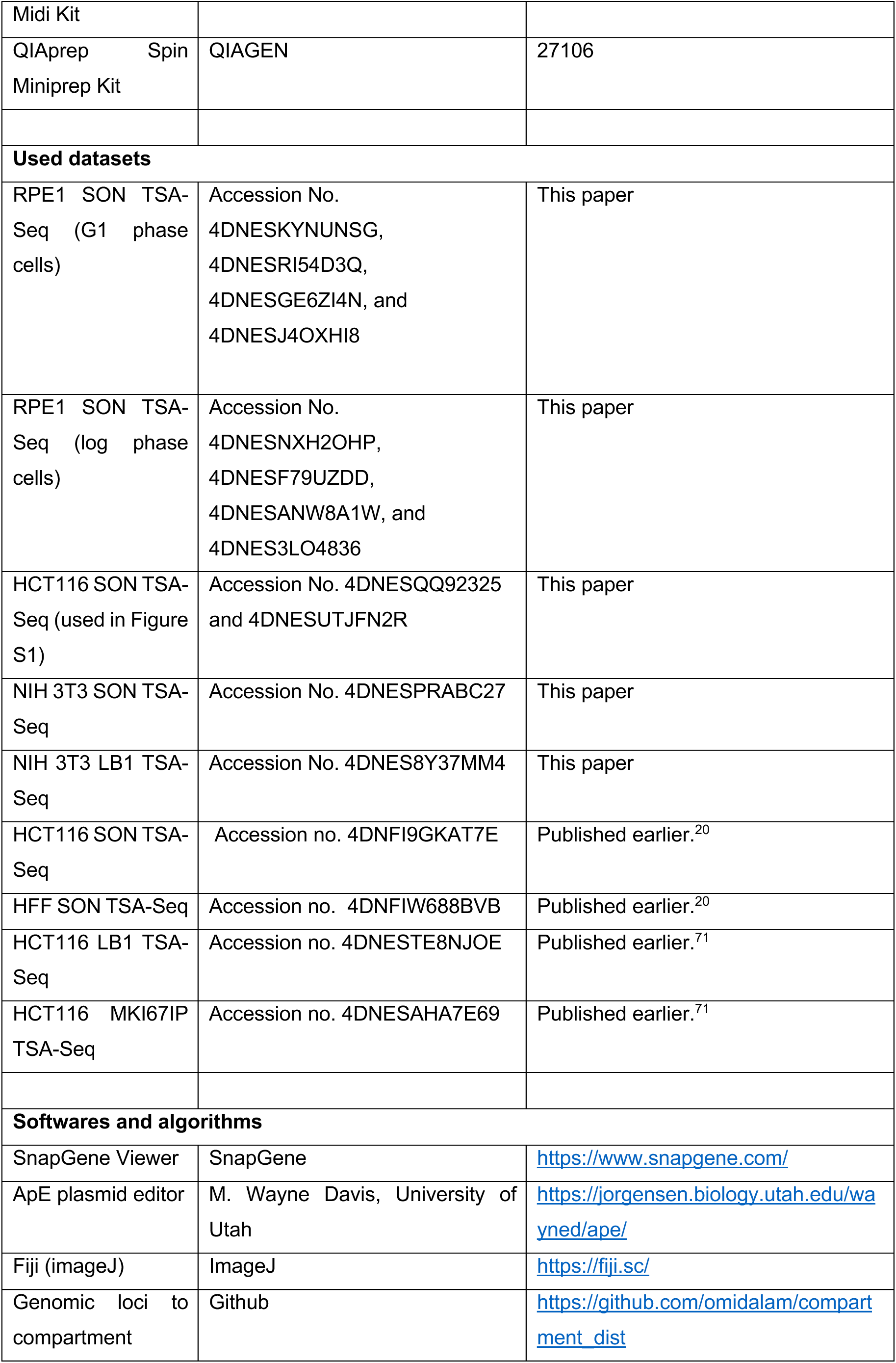
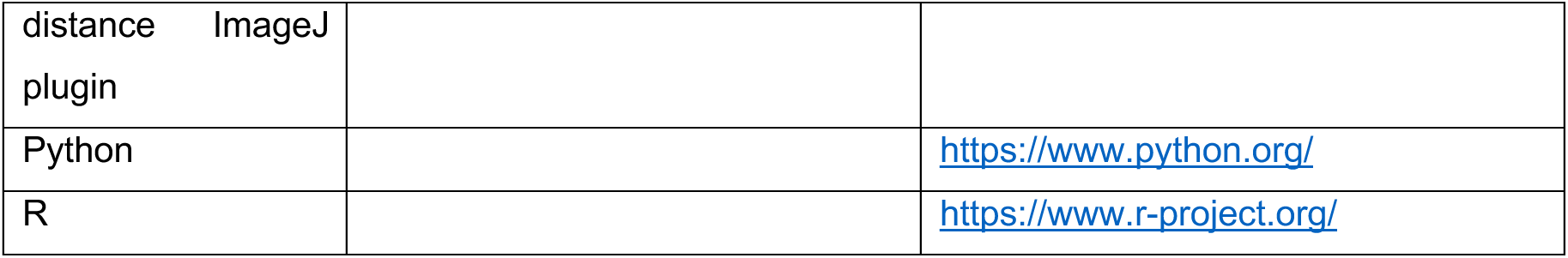

**• RESOURCE AVAILABILITY**

a. LEAD CONTACT

Further information and requests for resources and reagents should be directed to the corresponding author, Andrew Belmont.

**b. MATERIALS AVAILABILITY**

Details on antibodies/reagents used in the study have been included in the Key Resources Table. The reagents and constructs are available from the corresponding author upon reasonable request. Sequence details for all the constructs used in the study have been submitted to GenBank and corresponding accession numbers of constructs are provided in Table S1.

**c. DATA AND CODE AVAILABILITY**

All data used for plots in Figures and Supplementary Figures is supplied in Supplementary Data as Table S2. The Loci-Compartment Plugin for imageJ was used to measure distances relative to NS. The plugin is freely available and can be accessed at https://github.com/omidalam/compartment_dist.

**EXPERIMENTAL MODEL DETAILS**

**a. CELL LINES**

Mouse NIH 3T3 fibroblasts (ATCC CRL-1658) were grown in Dulbecco’s modified Eagle medium (DMEM with 4.5 g/l D-glucose, 4 mM L-glutamine, 1 mM sodium pyruvate and 3.7 g/l NaHCO3) supplemented with 10% HyClone Bovine Calf Serum (Millipore-Sigma, Cat # 12133C). HCT116, BJ-5ta-hTERT (ATCC CRL-4001), and RPE1-hTert cells were obtained through the ATCC and cultured according to the 4DN recommended culture conditions (https://4dnucleome.org/cell-lines/).

**• METHOD DETAILS**

a. **PLASMID DETAILS**

**Construction of retrofitting vector (pMini-HB-Tet96-attB-Neo-Puro-HSVTk-PB)**

The plasmid pMini-HB-attB-Neo-Puro-HSVtk-PB (hereafter referred to as the retrofitting construct, Ctrl V1.0) was assembled in a stepwise manner. First, an intermediate plasmid, pMini-HB, was constructed by ligating an 826 bp PCR amplicon (5HB-BAC-fw / 3HB-BAC-rev) derived from the DHFR-RSLB1 BAC,^34^ containing sequences homologous to common BAC vectors (e.g. pBeloBac11 and pBACe3.6) into the pMiniT 2.0 vector (NEB, E1203S). A Gibson Assembly (NEB) approach was then used to insert three PCR-amplified fragments into pMini-HB: (i) a 2 kb attB–promoterless Kan/Neo cassette (attB-Neo_fwd / attB-Neo_rev) from the attB-Neo-SacB plasmid, (ii) a 382 bp 5′ piggyBac (PB) ITR fragment (5PB_fwd / 5PB_rev) from pJoyC030, and (iii) a 3.4 kb vector backbone fragment (pMini-HB_fwd / pMini-HB_rev) from pMini-HB.

The resulting construct, pMini-HB-attB-Neo-5′PB, was PCR amplified using 5HB_KpnI-fw / 5PB_NheI-rev to generate a 5.8 kb amplicon, which was digested with DpnI, KpnI, and NheI, and ligated to a 6.6 kb KpnI/SpeI fragment from the plasmid pHSV-JT96-Puro. This second fragment contained the 3′ PB ITR, attB-promoterless Kan/Neo, SV40 promoter-driven PuroR, and EF1α promoter–driven HSVtk counterselectable marker. The pHSV-JT96-Puro plasmid was previously generated by inserting a 1.4 kb SV40 promoter-PuroR cassette into the SpeI site of pHSV-JT96-EFS-attP-Bla (Kowalczyk et al., manuscript in preparation). The attP sequence used in these constructs was obtained from the FRT-PGK-FRT-attP-Puro-DTA plasmid.

This retrofitting construct (Ctrl V1.0) was used for PB mobilization-based experiments. For *attP*-*attB* site-specific integrations, the retrofitting construct (Ctrl V1.0) was modified to a second-generation plasmid (pMini-HB-attB-Neo vector or Ctrl V2.0) through deletion of HSVtk-PB-Puro fragment. This deletion was achieved by digesting Ctrl V1.0 plasmid with SpeI and XbaI, followed by relegation of the 8.8 kb vector backbone. This modification of Ctrl V1.0 eliminates all the expressed eukaryotic cassettes, reducing the effect of integrated plasmid sequences on NS-targeting activity in our site-specific integration assays.

### Retrofitting of BAC vector for stable cell line creation

The vector backbone of selected BACs was retrofitted with fragments shown in Figure 2F through λ Red-mediated BAC recombineering as per the published protocols.^34,35,38^ A 10.0 kb retrofitting fragment with ∼ 410 bp homology ends for recombineering was prepared from Ctrl V1.0 plasmid through XhoI and PacI restriction digestion. *E. coli* strain SW102 was used for BAC recombineering. Recombinants containing the retrofitting cassette were selected for resistance to kanamycin (100 μg/ml) at 30°C on LB agar plates. The integrity of BAC constructs was verified by restriction enzyme fingerprinting.

### Construction of COL1A1 BAC derivative fragments

The construction of COL1A1 BAC fragments from COL1A1 BAC (RP11-267M22) involved several steps of BAC recombineering that enabled insertion or deletion at intended positions within the BAC. Initially, COL1A1 BAC intermediates F1, F2, and F3 were constructed by introducing the 3’PB-EF1α-BSD cassette from the template plasmid pJoyTet96lxpZeo-Bsd using primers specific to each derivative: 3PB-bsd-rec1-For and 3PB-bsd-rec1-newrev for F1, 3PB-bsd-rec2-For and 3PB-bsd-rec1-newrev for F2, and 3PB-bsd-rec3-For and 3PB-bsd-rec1-newrev for F3. Following this, the 5’PB-Tet96-Zeo cassette, derived from the same template by XbaI digestion, was subcloned into pMini-Col1A1-reg1 or pMini-Col1A1-reg2 at XbaI or NheI sites, respectively. The combinations for recombineering were as follows: COL1A1-F2 with pCol1A1reg1-5PB-Tet96 yielded COL1A1 F4 BAC; COL1A1-F3 with pCol1A1reg1-5PB-Tet96 yielded COL1A1 F5 BAC; COL1A1-F2 with pCol1A1reg2-5PB-Tet96 yielded COL1A1 F6 BAC; COL1A1-F3 with pCol1A1reg2-5PB-Tet96 yielded COL1A1 F7 BAC; COL1A1-F1 with pCol1A1reg1-5PB-Tet96 yielded COL1A1 F8 BAC; and COL1A1-F1 with pCol1A1reg2-5PB-Tet96 yielded COL1A1 F9 BAC. Further modifications through recombineering in BAC vector backbone were made to the F7 and F8 BACs by inserting a promoterless-Neo and attB fragment, resulting in COL1A1-F7attB and COL1A1-F8attB with kanamycin resistance.

The next phase involved Galk-based recombineering where F10attB was constructed by removing a 29.6 kb region from F7attB (Primers: recColD7GalkFor/ recColD7GalkRev; template pUGG plasmid; product size 1.3 kb). This F10attB BAC served as intermediate to create subsequent BACs F11 through F15. The primer combinations for GalK-based recombineering were as follows: COL1A1-F11 (recD11-D12fw/ recD11rev); COL1A1-F12 (recD11-D12fw/recD12-D15rev); COL1A1-F13 (recD13fw/ recD13-D14rev); COL1A1-F14 (recD14-15fw/recD13-D14rev); and COL1A1-F15 (recD14-15fw/recD12-D15rev). Galk-based 99 kb deletion in COL1A1 F8 BAC was made to derive COL1A1 F16 BAC (Primers recColD8GalkFor/ recColD8GalkRev; Template pUGG plasmid).

The intergenic construct COL1A1 F17 was made by ligating two PCR-amplified fragments derived from F13 BAC into the pMini2.0 vector. A 4.6 kb fragment from first intermediate construct (pMini-COL1A1 prod 1) was excised by BamHI and ligated to second intermediate construct (pMini-COL1A1 prod 2) at unique BamHI site. The final construct was verified by PCR and restriction digestion for correct orientation of insert. This 11 kb intergenic COL1A1-SGCA fragment was cut using PmeI/ZraI restriction enzymes and subcloned to PmeI digested retrofitting vector Ctrl V1.0 vector to get COL1A1-F17-RFTD construct.

### COL1A1 F17 random deletion library

A random fragment library of a minimal speckle target region of COL1A1 BAC was prepared by excising and eluting a 11 kb fragment from COL1A1-F17-RFTD using PacI//PmeI/SpeI-HF (NEB). About 6 μg of the eluted linear plasmid was fragmented to 1000-9000 bp using a Bioruptor Pico (Diagenode) machine with a mode of 7 sec ON – 30 sec OFF for 1 sonication cycle. The sonicated DNA was eluted on agarose gel to retrieve fragments ranging from 2-8 kb, end repaired (End-It DNA End-Repair Kit: Epicentre Biotech ER0720), and blunt-end ligated to Ctrl V2.0 vector. The plasmid was digested with XbaI leaving behind a minimal vector containing-96mer, *attP*-Promoterless Kan/Neo cassette, followed by blunting by end-filling with Klenow (M0210, NEB) at 25°C. The ligated mix was transformed in NEB 5-alpha Competent *E. coli* (C2987H). Plasmids derived from randomly selected colonies were digested, and end sequenced to verify the inserts. The self-ligated vector without insert was used as the corresponding “empty vector” (Ctrl V2.0) negative control plasmid for experiments screening the fragment library.

### Cloning of SPAD fragments

HCT116 template gDNA was used to PCR amplify fragments spanning H3K27ac peaks at chr12:6,531,445-6,536,855 (5411 bp), chr20:63,656,173-63,660,806 (4634 bp), and chr6:31,813,579-31,818,081 (4503 bp) using primers 369N23-D03-Fw/369N23-D03-Rev, 3184A7-D01-Fw/3184A7-D01-Rev, and D01_92G8_Fw/D01_92G8_Rev, respectively. The PCR products were gel purified and ligated to linearized Ctrl V1.0 vector.

### Construction of TetR-TF fusion plasmids

Second generation lentiviral constructs expressing TetR-EGFP fused to transcription factors of interest were created by using Gibson Assembly approach (NEB). Briefly, the lenti F9-TetR-EGFP-IRES-Pur vector published elsewhere,^72^ was linearized by BsrGI-HF restriction endonuclease (NEB), gel eluted and assembled with PCR amplified transcription factor CDS (coding sequence) as per manufacturers’ instructions. The CDS for TFAP2A, MXI1, ZBTB18, TWIST1, ZNF384 were amplified from cDNA prepared from U2OS cells (First strand synthesis kit, ThermoFisher Scientific), cloned in pMini2.0 vector (NEB), and verified by Sanger’s sequencing. The verified CDS were PCR amplified using indicated primers, incorporating a polyG-S (Glycine-Serine) linker to enable flexibility between TetR-EGFP and TF fusion protein. The codon-optimized ZNF460 CDS was synthesized by ThermoFisher, while CDS for TFAP2B (Cat no. DQ894621) and TFAP2C (Cat no. BC035664) were procured from Transomic technologies, Inc (Huntsville, AL, USA). The lentiviral constructs expressing TetR-EGFP fused AADs were generated by subcloning AAD fragments derived from full-length TFs mentioned above or AAD constructs described earlier.^52^ The AAD fragments were PCR amplified and subcloned between BsrGI/PshAI sites of lenti tetR-EGFP-IRES-Puro plasmid and verified by whole plasmid sequencing.

### Mutational screen for COL1A1 D45 fragment

The predicted transcription factor binding sites on D45 fragment (628 bp) were eliminated in a synthetic fragment (F45-allmut), ordered through Integrated DNA Technologies (IDT). The product was PCR amplified (45frgE-fw/45frgA-Rev) and cloned into the PmeI/NdeI digested and dephosphorylated RFTDv2.0 vector (8.6 kb). The transcription factor binding sites in this construct were separately restored to the original individual or overlapping TF binding sites through site-directed mutagenesis. The primers used to create the final constructs (RFTDv2.0-D45-FragF rSDM1/2/3/4/5) are listed in the Table S1.

**b. DRUG TREATMENTS**

**TRANSCRIPTIONAL INHIBITION**

Transcriptional inhibition in RPE1 cells was performed for 4 hours on cells in log phase growth at 70% – 80% confluency with final concentrations of 0.1 μg/mL (278 nM) Triptolide, or 50 μg/mL (157 μM) DRB, or DMSO as the solvent-only control. In each experiment, we adjusted the volume of DMSO to make sure all groups of cells were exposed to the same DMSO concentration.

RPE1 cell synchronization was achieved by dual mitotic shake-off of cells. At ∼80–90% confluency the asynchronous cells were subjected to a first mitotic shake-off to remove cells already in mitosis. The flasks were then rinsed with fresh media and treated with 50 ng/mL nocodazole in fresh media for 12 hours to arrest cells in mitosis. After 12 hours of incubation, we shook off the arrested mitotic cells (∼4 million shaken off from one T300 flask) and transferred the cells within the nocodazole-containing media into new T150 flasks and added either 0.1 μg/mL Triptolide, DMSO, or nothing (for the “untreated” sample). The cells were incubated for an additional 1 hour for a combined mitotic and transcriptional arrest (the arrested cells won’t attach). After 1 hour of combined mitotic and transcriptional arrest, the cells were harvested by centrifugation at 200x*g*, and rinsed twice with fresh media containing triptolide (or DMSO or untreated control) to wash away nocodazole. The cells released from mitotic arrest into G1 phase were plated into fibronectin-coated T150 flasks and cultured in fresh media containing Triptolide (or DMSO) for 3 hours prior to cell harvesting. To coat flasks, fibronectin was diluted 1:100 in HBSS and added to flasks (15 mL for a T150 flask) and incubated at room temperature (RT) for 1 hour in cell culture hood immediately before use. After coating of the flasks, the fibronectin/HBSS was removed and cells/media were immediately added into the flasks.

**EU-PULSE LABELING:** RPE1 cells were grown on glass coverslips with or without transcription inhibitors as described above. 5-Ethynyl uridine (50 mM stocks in 50% DMSO/ 50% PBS) was added to media at 1mM final concentration and pulse-labelled for 20 mins. After EU labeling the cells were rinsed with PBS three times, fixed with freshly made 3.7% paraformaldehyde at room temperature (RT) for 12 mins, washed with PBS for 3 x 5 mins, and then permeabilized with 0.5% triton X-100 /1X PBS (PBST) at RT for 15 mins. EU was Click-it labeled with Alexa 647-azide as previously described.^73^

**c. GENERATION OF LENTIVIRAL PARTICLES**

Lentiviral particles were produced using the packaging plasmid pCMV-dR8.2 (Addgene plasmid #8455) and the envelope plasmid pCMV-VSV-G (Addgene plasmid #8454).^74^ HEK293T cells were seeded in a 12-well plate and cultured to 60–80% confluency prior to transfection. For each lentiviral preparation, cells were cotransfected with 500 ng of the lentiviral vector, 450 ng of pCMV-dR8.2, and 50 ng of pCMV-VSV-G using lipofectamine 2000, following the manufacturer’s protocol. After 24 hours, the culture medium was replaced with fresh DMEM supplemented with 10% fetal bovine serum (FBS). Seventy-two hours post-transfection, viral supernatants were harvested and filtered through a 0.45 μm syringe filter to remove cell debris. All transductions were performed at a low multiplicity of infection (MOI). For transduction, 0.5-1 × 10⁵ cells were seeded per well in a 12-well plate and incubated with viral supernatant in the presence of 8 μg/mL polybrene for 24 hours. After 24 hours, the media was replaced with culture media supplemented with 1.0 μg/mL (NIH 3T3) or 1.2 μg/mL (HCT116) puromycin and maintained for another 4-5 days. For TetR-EGFP fusion protein tethering assays, NIH 3T3 cells were transduced with lentiviral particles expressing TetR–EGFP fusion proteins and selected with 1.0 µg/mL puromycin for 5 days prior to fixation.

**d. GENERATION OF STABLE AND KI LINES**

**ESTABLISHMENT OF PB MOBILIZED CELL LINES:** BAC DNA for transfection of mammalian cells was prepared with the QIAGEN Large Construct Kit as per the manufacturer’s instructions. All other plasmid constructs for transfection were purified by Qiagen spin miniprep or midiprep kits. Lipofectamine 2000 was used to transfect the cells with BAC or plasmid DNA according to the manufacturer’s instructions. PB transposition of BACs was done by cotransfection of 10 μg BAC and 3 μg of inducible PB transposase expressing plasmid (mPB-L3-ERT2-tatRRR-mCherry). Nuclear translocation of the expressed PB transposase was achieved by supplementing the culture media with 2μM tamoxifen for 48 hours. The transfected cells were enriched in the appropriate culture medium supplemented by Puromycin for two weeks of puromycin selection, followed by 5 days of 10 μg/ml ganciclovir counterselection. Mixed clonal populations of stable transformants were expanded and either transduced with lentiviruses or transfected with plasmids expressing TetR-EGFP for 4-5 days, as described in the preceding Section. Individual cell clones were obtained by serial dilution or retrieval of colonies using filter discs.^75^

**TARGETED KNOCK-IN OF NIH 3T3 CELLS**: NIH 3T3 cells were engineered to harbor a docking cassette comprising an attP site, a 96mer TetO array, and an EFS promoter-driven HSV-tk-IRES-BSD cassette at the indicated LAD or iLAD genomic loci (Figure S3). Genomic cleavage at the target sites – Reg 01 (Chr1:19564883–19568306, mm10) and Reg 03 (Chr1:105892227–105892428, mm10) was achieved by Cas9 and guide RNAs (gRNAs) expressed from the pX330A-1x3 vector (Addgene plasmid #58767).^76^ For each locus, two distinct gRNA constructs were designed. The gRNA oligonucleotides were cloned into the BbsI-digested pX330A-1x3 backbone via Golden Gate assembly as per the recommended protocol.^76^

Donor plasmids for homology-directed repair were constructed by amplifying ∼2 kb genomic fragments-chosen to contain unique AgeI and SacII restriction sites towards the center of the fragments-from wild-type NIH 3T3 DNA using the following primer pairs: LAD1Fw/LAD1Rev (Reg 01) and LAD3Fw/LAD3Rev (Reg 03). Amplicons were cloned into the pMini2.0 vector (NEB PCR cloning kit). These plasmids were then used as templates to generate linear donor fragments via PCR, designed to remove gRNA target sites while retaining 800–900 bp homology arms on either side. Primer pairs used for generating linearized donor fragments were HB_LAD1_NFw/HB_LAD1_Rev and HB_LAD3_NFw/HB_LAD3_Nrev.

A donor cassette (∼6.3 kb), including the attP-TetO-HSV-tk-IRES-BSD sequence, was PCR-amplified from the pDTA-J-EFS-attP-HSVtk-IRES-BSD-Tet96 plasmid using primers EFSattP_Fw and EFS_attP_Rev. The cassette was ligated into the homology arm-containing vectors via AgeI and SacII restriction sites to generate final donor constructs (LAD1-donor and LAD3-donor). These final donor constructs were linearized with NotI and PmeI, gel-purified, and co-transfected with their respective gRNA/Cas9 plasmids into NIH 3T3 cells at a 1:2 molar ratio (donor:gRNA plasmid).

**SITE-SPECIFIC RECOMBINATION BY ΦC31 INTEGRASE:** Site-specific integration of specific DNA fragments at the intended locus within NIH 3T3 cells was achieved through the recombination of *attB* on BACs/plasmids and target *attP* sites previously integrated in the docking lines.^72^ The φC31 integrase, derived from Streptomyces phage φC31, is a site-specific recombinase that can efficiently recombine *attP* and *attB* sequences.^39,40^ The *attB* site on BACs/plasmids contain a promoterless Neo^R^ cassette positioned such that precise insertion at the *attP* docking site replaces the Blasticidin resistance cassette (BSD) from the docking target site with the Neo^R^ cassette, leading to expression of the neomycin resistance gene via the constitutive elongation factor 1 alpha (EF1α) promoter (Figure 3D).

To achieve integration of BACs/plasmids at either the iLAD (Reg 03) or LAD (Reg 01) docking sites, the BACs/plasmids and φC31 integrase expression vector plasmid (pCAG integrase) were cotransfected at a 1:2 molar ratio (BAC/plasmid:pCAG integrase). After 48 hours of transfection, the transfected NIH 3T3 cells were enriched in DMEM culture medium supplemented by G418 for three weeks, followed by 5 days of 10 μg/ml ganciclovir counterselection. Individual cell clones were obtained by retrieval of colonies using filter discs,^75^ followed by transient expression of TetR-EGFP through lentiviral transduction for 4-5 days.

**e. IMMUNOFLUORESCENCE**

Cells grown on glass coverslips were fixed in freshly prepared 2% paraformaldehyde (PFA) in 1XPBS buffer for 15 min at room temperature (RT). Cells were then washed with PBS for 2 x 5mins and with 0.1% PBST for 5 mins. Cells were blocked with 5% normal goat serum at RT for 1 hour and then incubated with rabbit anti-SON polyclonal antibody diluted 1:2000 and mouse anti-RNA Pol II Ser5P monoclonal antibody diluted 1:100 in blocking buffer at 4°C for 10-12 hours. Cells were washed with 0.1% PBST for 3 x 5 mins and then incubated with goat anti-rabbit-FITC and goat anti-mouse-Alexa594. Cells were washed with 0.1% PBST for 3 x 5 mins and then mounted with Mowiol-DABCO anti-fade media containing 0.3 μg/ml DAPI.^77^

**f. FLUORESCENCE IN SITU HYBRIDIZATION (FISH)**

**DNA FISH probes:** GAPDH, COL1A1, Dhfr/Msh3, and COL1A2 BACs were used to prepare probes for 3D DNA FISH. A biotinylated oligonucleotide (see Table S1) synthesized at IDT was used as a probe for FISH based detection of the TetO array. Preparation of biotin or digoxigenin labeled DNA FISH probes and 3D DNA FISH of interphase nuclei were carried out as described earlier,^18^ with small modifications. Instead of culturing cells on poly-L-lysine coated coverslips for 15 minutes, the cells were grown on untreated glass coverslips (12 mm diameter) for 3-4 days before fixation with 2% paraformaldehyde in 1XPBS for 15 mins at RT. FISH signals were detected by incubation with Alexa Fluor 647 conjugated Streptavidin (1:200 dilution) or Alexa 594 conjugated Streptavidin (1:200 dilution) for biotin-labeled probes, or Alexa Fluor 647 conjugated IgG fraction monoclonal mouse anti-digoxin (1:200 dilution) for digoxigenin labeled probes, diluted in SSCT with 1% Bovine Serum Albumin, for 2 hours at RT or overnight at 4°C. Coverslips were washed in SSCT for 4 × 5 min, rinsed with 4x SSC and mounted. All samples were mounted with a Mowiol-DABCO anti-fade medium containing ∼0.3 μg/ml DAPI.^77^

**g. TSA-SEQ**

SON TSA-Seq 2.0 procedure with enhancement condition E, reagents used, and data analysis protocols has been described previously.^20^ Specific conditions used for these data sets, such as antibody concentrations and source, numbers of cells, library construction, etc, are provided in the metadata for each dataset. The datasets and corresponding metadata sheets are publicly available at the 4D Nucleome server with accession numbers provided in the Key resources table.

**• QUANTIFICATION AND STATISTICAL ANALYSIS**

**a. IMAGE ACQUISITION AND ANALYSIS**

All images were acquired as 3D optical-section images with 0.2 μm z-steps using a V4 OMX (GE healthcare) microscope equipped with a 100X, 1.4 NA oil immersion objective (Olympus) and two Evolve EMCCDs (Photometrics). Image stacks were subjected to 10 cycles of constrained iterative deconvolution (conservative Ratio method) to remove out of focus blur, and corrected for camera and chromatic aberration misalignments using OMX image alignment by local triangulation method using SoftWoRx software (version 7.0) (GE Healthcare). All subsequent image analysis and preparation were done using Fiji software (ImageJ). Representative images were assembled using Illustrator (Adobe).

Sum intensity measurement for Figure 1 was made in FIJI on non-deconvolved images where the total intensity of 26 slices (0.2 μm in z-axis) was normalized to cytoplasmic background, exposure time, and % transmitted exciting light. The quantification methods were adapted from a previous publication.^78^

The SON antibody used in our study stains the core of NS,^9,15^ while the periphery of NS composed of U1, U2 snRNPs extends to an additional 20-50 nm beyond the NS core.^9^ Earlier the gene was considered NS associated when the gene was in physical contact with NS as observed with the fluorescence microscopy.^17,79^ However, accommodating the resolution of limits of fluorescence imaging we used a cut-off for NS-association when the transgene signal was detected within 0.25 μm of NS periphery. To ensure reproducibility of measurements and to negate the effects of background staining as well as the variable intensities of different NS, we defined the edge of the NS by where the pixel intensity fell to 40% of the maximum SON staining intensity of that NS. Distances relative to the nearest edge of the nearest NS were measured by using the Loci-Compartment Plugin for imageJ. The plugin is freely available and can be accessed at https://github.com/omidalam/compartment_dist.

**b. STATISTICAL ANALYSIS AND REPRESENTATION**

A z-test for two population proportions was used to assess statistical differences in the proportion of events within the 0–0.25 μm bin of the stacked histograms. The test was performed under the null hypothesis that the proportions in the two groups are equal, and corresponding p-values were calculated accordingly. The null hypothesis was tested using this equation:

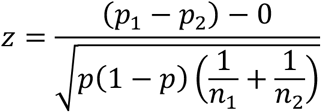

Here *p*_1_ and *p*_2_ are the sample proportions from the two groups, *n*_1_ and *n*_2_ are the sample sizes of the two groups, and *p* is the pooled proportion calculated as:

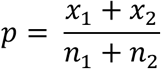

*x*_1_ and *x*_2_ are total counts in the bin (0-0.25 μm) from the two groups.

Statistical significance of differences in mean distances between samples was assessed using one-way ANOVA followed by a post hoc Tukey’s Honestly Significant Difference (HSD) test. P-values are represented in the figures as follows: n.s., not significant; * p < 0.05; ** p < 0.01; *** p < 0.001; **** p < 0.0001. Unless otherwise specified, data are presented as mean ± SEM (standard error of the mean). Statistical analyses were performed using Microsoft Excel, the R package ’stats’ (for ANOVA and HSD tests), and the online tool at https://www.socscistatistics.com.

### Data description

The TSA-Seq data generated in this study have been submitted to the 4DN Data Portal (https://data.4dnucleome.org/). The corresponding accession numbers can be found in the key resources table.

## ACKNOWLDEGMENTS

This work was supported by the National Institute of General Medical Sciences grant R01GM058460 (A.S.B) and the National Institutes of Health Common Fund 4D Nucleome Program grants U54DK107965 (A.S.B.,H.Z.) and UM1HG011593 (A.S.B). We thank Bas van Steensel (The Netherlands Cancer Institute, the Netherlands) for providing pJoyC030 and mPB-L3-ERT2-tatRRR-mCherry plasmids, and David Gilbert (San Diego Biomedical Research Institute, USA) for FRT-PGK-FRT-attP-Puro-DTA and attB-Neo-SacB plasmids. We also acknowledge contribution of the Flow Cytometry Facility and the DNA Services Laboratory, Roy J. Carver Biotechnology Center, University of Illinois Urbana-Champaign for FACS sorting and NGS sequencing, respectively. We thank Dr. Sandra Kay McMasters for providing cell culture media (UIUC, SCS Cell Media Facility).

## AUTHOR CONTRIBUTIONS

P.C. and A.S.B conceived of and designed the study; P.C. performed most experiments including TSA-Seq in HCT116 cells, DNA-FISH, stable cell line generation, BAC dissection, imaging, data analysis and interpretation. P.G. identified NS-targeting of well-characterized AADs and contributed to performing and analyzing TetO-TetR tethering experiments. L.Z. conducted TSA-Seq experiments in RPE1 cells and performed data analysis and interpretation for RPE1 TSA-seq. M.Z. and H.Z. were instrumental in establishing the site-specific integration system in HCT116 cells, which was modified further and used in NIH 3T3 cells by P.C. P.C. and A.S.B. wrote the manuscript with input from other authors.

## Supplementary Figure Legends

**Figure S1. Genome organization relative to NS is maintained independent of transcription in log phase HCT116 cells.**

**(A)** Genome browser view of SON TSA-Seq enrichment profiles in untreated, DMSO-, or TPL-treated log phase HCT116 cells.

**(B-C)** Two-dimensional histograms showing correlations of smoothed SON TSA-Seq enrichment scores between biological replicates of DMSO treatment **(B)** or DMSO versus TPL treatment **(C)**. Colors indicate the number of 20 kb genomic bins with each 2D TSA-Seq score interval (∼0.02 X 0.02). Pearson’s r denotes the Pearson product-moment correlation coefficient.

**Figure S2. Transcription-independent NS-association of endogenous genomic loci in HCT116 cells.**

**(A)** Genome browser view of GAPDH, COL1A1, and COL1A2 loci in the indicated cell lines showing proximity to NS measured by TSA-Seq SON (top three tracks) and expression levels measured by RNA-seq scores (bottom three tracks).

**(B)** Maximum intensity z-projections of representative 3D DNA FISH images (red) of endogenous GAPDH, COL1A1, and COL1A2 loci in HCT116 cells after DMSO or DRB treatment for 4 hours. Cells were immunostained for NS (green) and counterstained with DAPI (blue). Scale bars, 2 μm (main panels) and 0.5 μm (insets).

**(C)** Box plots showing distances of FISH-labeled genomic loci to the nearest NS in HCT116 cells. Plots indicate median (black horizontal line), mean (black square), individual data points (gray diamonds), and 1.5X interquartile range (whiskers). Statistical significance of DRB treatment was compared with respective DMSO controls (n = 162-169). Significance was tested by one-way ANOVA with post hoc Tukey’s HSD. n.s., not significant at P < 0.05.

**(D)** Schematic of PiggyBac (PB) transposition approach for candidate BAC screen in HCT116 cells.

**Figure S3. Generation of NIH 3T3 docking lines for φC31-mediated site-specific integration.**

**(A)** Chromosome-wide Lamin B1 and SON TSA-Seq profiles in mouse cells highlighting regions (red) selected for CRISPR-Cas9 based insertion of an attP site, TetO-96mer array, and an EFS promoter-driven HSVtK-IRES-BSD cassette.

**(B)** Maximum-intensity projections of two optical z-sections from representative nuclei of independent clones showing the position of mobilized TetO arrays (green) relative to NS (SON, red). DNA was counterstained with DAPI (blue). Scale bars, 5 μm.

**(C)** Stacked histograms showing distance of TetO-96mer signals to the nearest NS in the indicated NIH 3T3 knock-in clones. x-axis: construct; y-axis: percentage of foci within each distance bin. (n=87-99).

**(D)** Box plots showing distances of TetO foci from NS in the indicated NIH 3T3 knock-in clones. Plots indicate median (black horizontal line), mean (black square), individual data points (gray diamonds), and 1.5X interquartile range (whiskers) (n=87-99).

**Figure S4. Predicted transcription factor binding motifs within the 600 bp COL1A1 fragment F45.** Sequence logos of transcription factor binding motifs predicted using the human JASPAR 2022 database. The x-axis indicates the relative position within the consensus binding site, and the y-axis shows the information content (bits), reflecting the degree of nucleotide conservation. The height of each nucleotide (A, C, G, T) within a stack represents both its frequency and contribution to binding specificity. Below each motif, the corresponding 600 bp wild-type (WT) sequence and its mutant version are shown, with mutated bases are highlighted in red and altered motif shaded in gray.

**Figure S5. Predicted acidic activation domains (AADs) in the indicated transcription factors identified with ADpred.**

**(A)** Graphical representations of predicted AADs within the indicated transcription factors, generated using ADpred with default settings. The x-axis shows the relative position within the full-length protein sequence (scaled by amino acid number), and the y-axis shows the probability of activation domain function. The red horizontal line marks the prediction threshold. Blue plots indicate AADs within acidic TFs, red plots indicate AADs within other TFs enriched in non-acidic activation domains (NAADs) or CTCF protein. Gray shaded regions mark peptide sequences selected for functional testing in the tethering assays.

**(B)** Summary of peptide sequences tested in the tethering assay. For each entry, the protein name, UniProt ID (canonical isoform), peptide sequence, and its position within the reference protein are listed. Acidic residues (D, E) are colored green, hydrophobic residues (F, M, I, L, V, C, W) in red, and basic residues (K, R) in blue.

## Supplementary Table Legends

Table S1. Reagent information (Plasmid/BAC Constructs and Oligonucleotides) used in this study.

Table S2. All data and statistical analysis used to prepare plots in Figures and Supplementary Figures of the study.

