## Supplementary Figures for "Acidic transcription factors position the genome at nuclear speckles through transcription dependent and independent mechanisms"

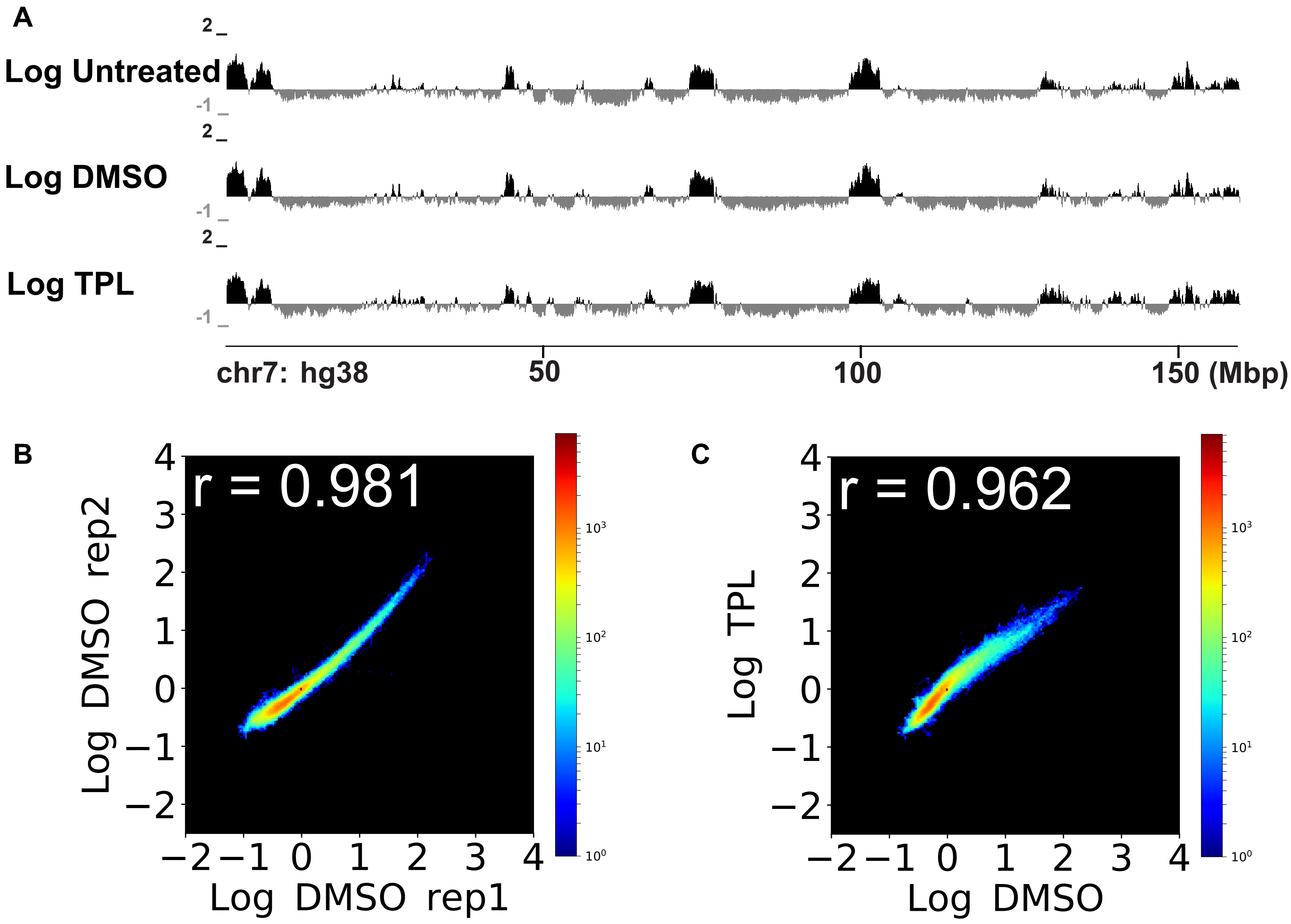
A
Log Untreated
Log DMSO
Log TPL
2
-1
chr7: hg38
50
100
150 (Mbp)
B
r = 0.981
Log DMSO rep2
Log DMSO rep1
C
r = 0.962
Log TPL
Log DMSO

**Figure S1**

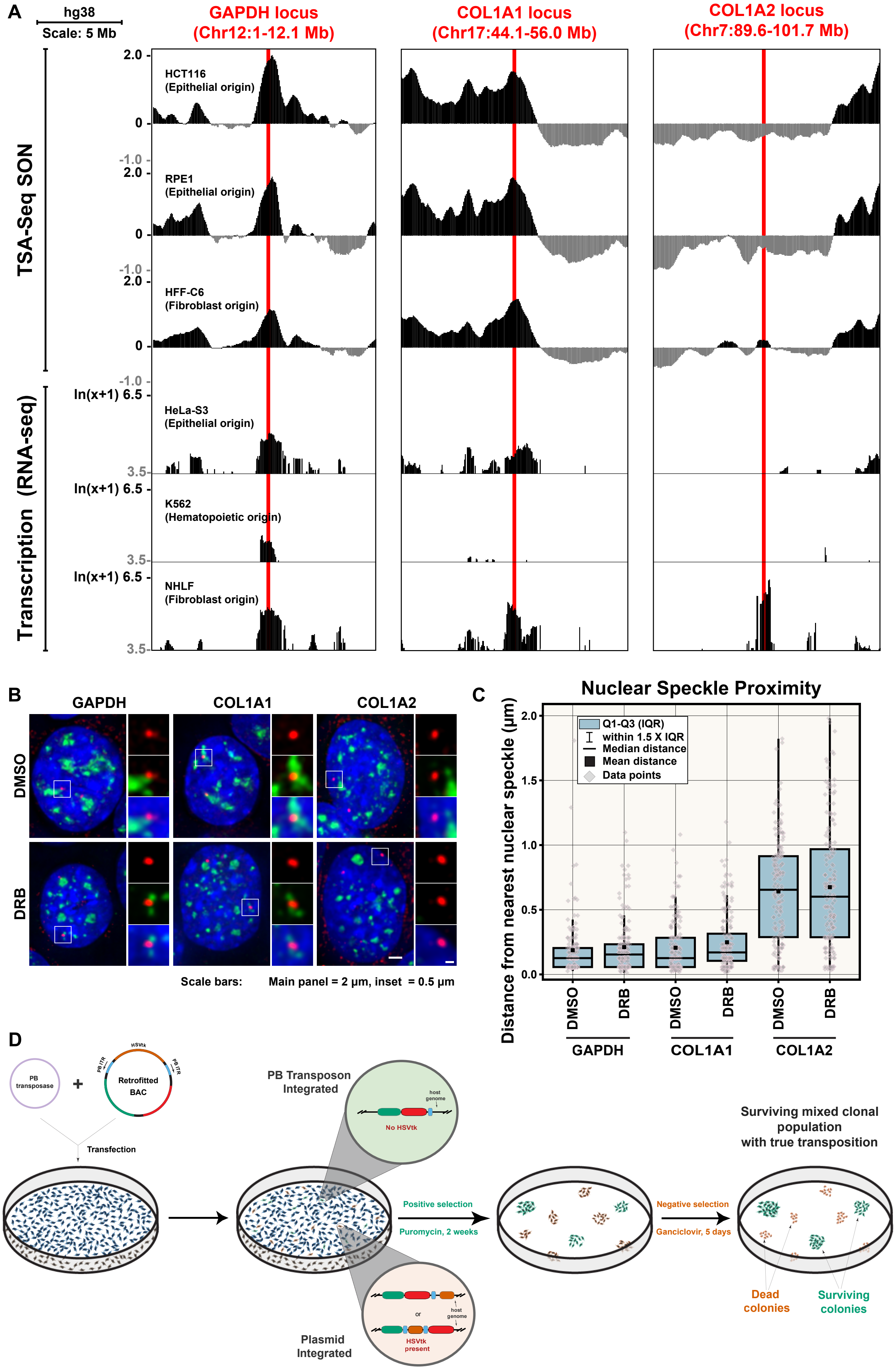
A
hg38
Scale: 5 Mb
GAPDH locus (Chr12:1-12.1 Mb)
COL1A1 locus (Chr17:44.1-56.0 Mb)
COL1A2 locus (Chr7:89.6-101.7 Mb)
TSA-Seq SON
HCT116 (Epithelial origin)
RPE1 (Epithelial origin)
HFF-C6 (Fibroblast origin)
2.0
0
-1.0
Transcription (RNA-seq)
ln(x+1) 6.5
3.5
HeLa-S3 (Epithelial origin)
K562 (Hematopoietic origin)
NHLF (Fibroblast origin)
B
GAPDH
COL1A1
COL1A2
DMSO
DRB
Scale bars: Main panel = 2 μm, inset = 0.5 μm
C
Nuclear Speckle Proximity
Distance from nearest nuclear speckle (μm)
Q1-Q3 (IQR)
within 1.5 X IQR
Median distance
Mean distance
Data points
0.0
0.5
1.0
1.5
2.0
DMSO
DRB
GAPDH
COL1A1
COL1A2
D
PB transposase
Retrofitted BAC
HSVtk
PB ITR
Transfection
PB Transposon Integrated
host genome
No HSVtk
Plasmid Integrated
or
HSVtk present
Positive selection
Puromycin, 2 weeks
Negative selection
Ganciclovir, 5 days
Surviving mixed clonal population with true transposition
Dead colonies
Surviving colonies

### Figure S2

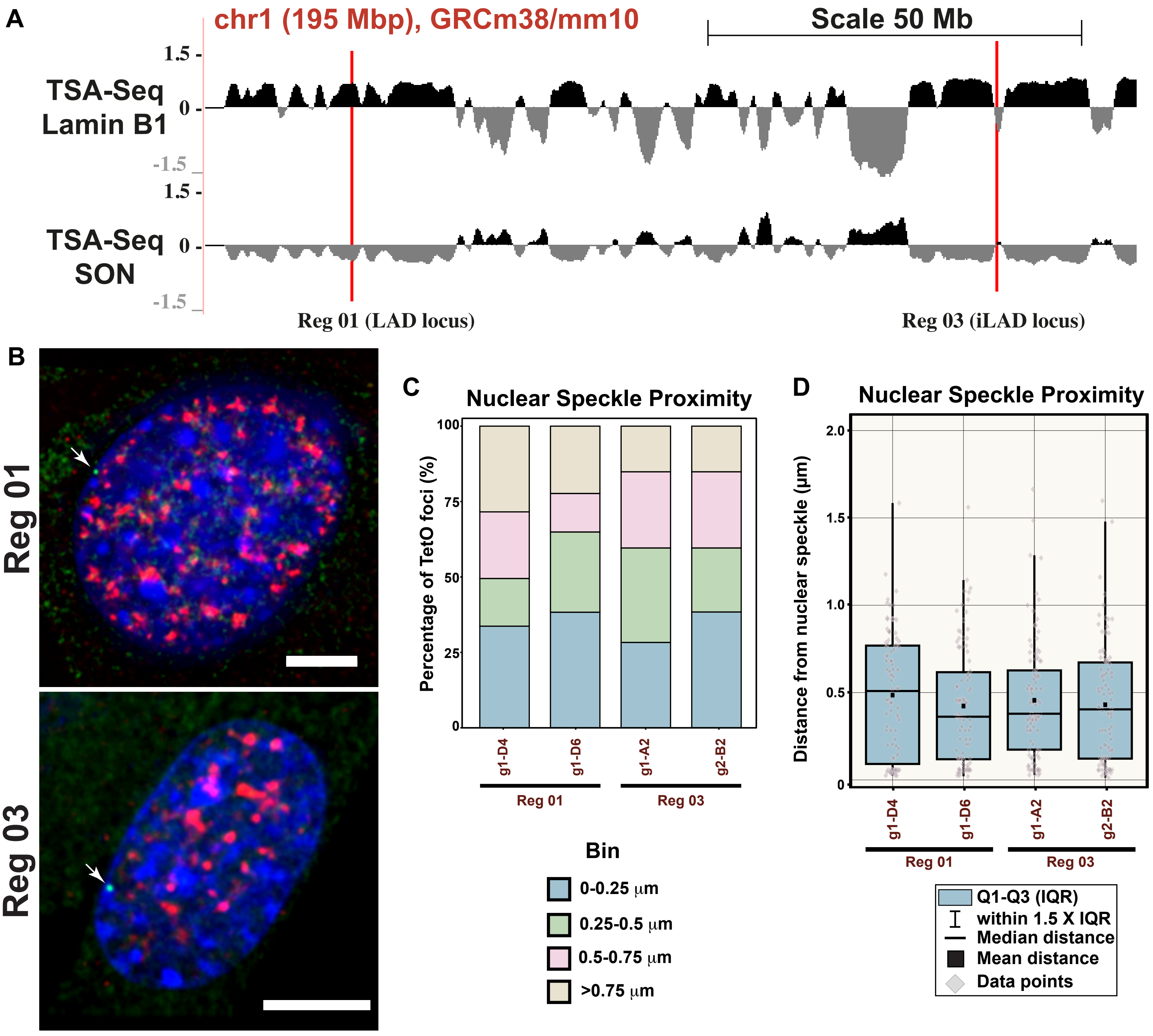

A
chr1 (195 Mbp), GRCm38/mm10
Scale 50 Mb
TSA-Seq
Lamin B1
1.5
0
-1.5
TSA-Seq
SON
Reg 01 (LAD locus)
Reg 03 (iLAD locus)
B
Reg 01
Reg 03
C
Nuclear Speckle Proximity
Percentage of TetO foci (%)
100
75
50
25
0
g1-D4
g1-D6
g1-A2
g2-B2
Reg 01
Reg 03
Bin
0-0.25 μm
0.25-0.5 μm
0.5-0.75 μm
>0.75 μm
D
Nuclear Speckle Proximity
Distance from nuclear speckle (μm)
2.0
1.5
1.0
0.5
0
g1-D4
g1-D6
g1-A2
g2-B2
Reg 01
Reg 03
Q1-Q3 (IQR)
within 1.5 X IQR
Median distance
Mean distance
Data points


### Figure S3

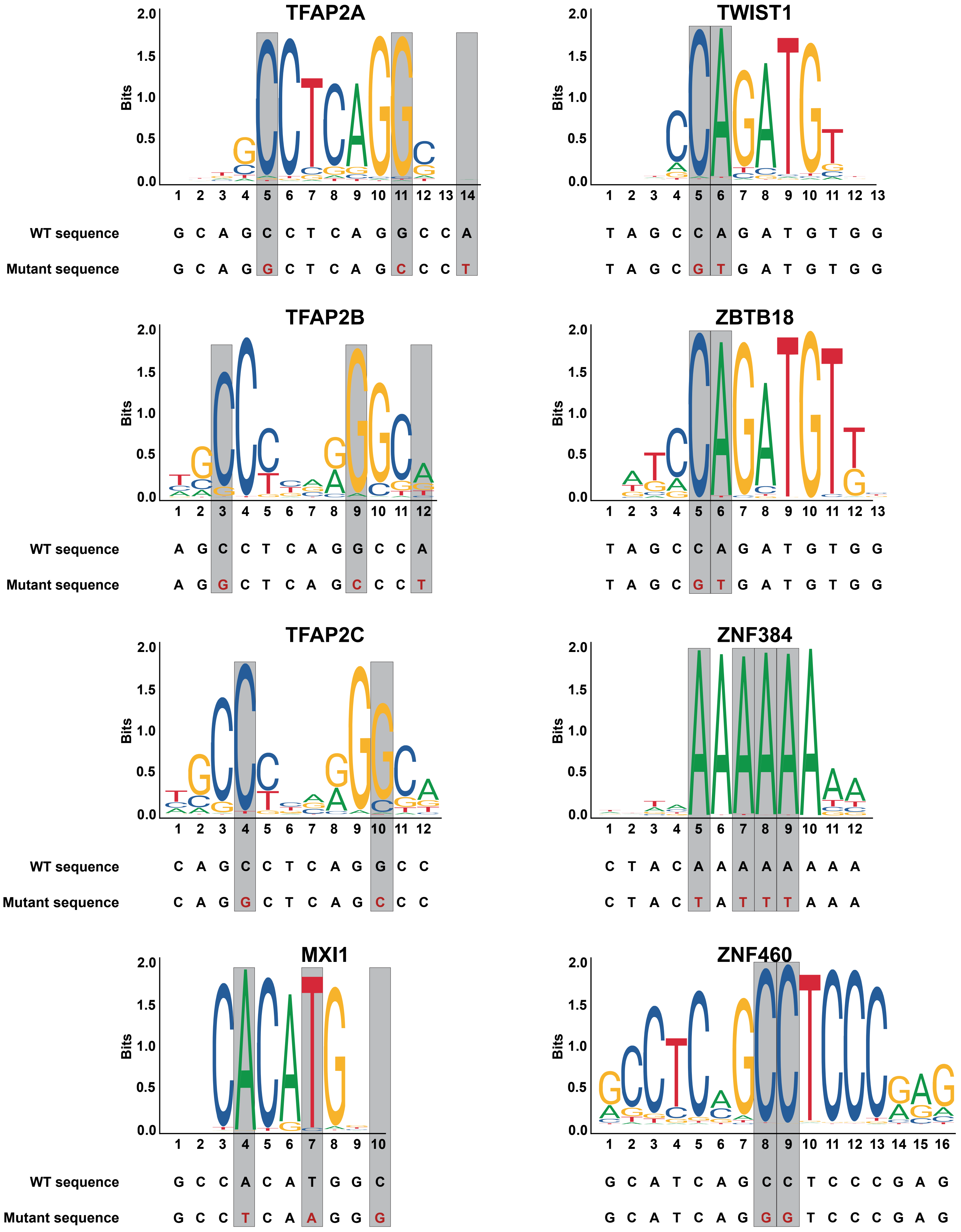

TFAP2A
WT sequence G C A G C C T C A G G C C A
Mutant sequence G C A G G C T C A G C C C T
TWIST1
T A G C C A G A T G T G G
T A G C G T G A T G T G G
TFAP2B
WT sequence A G C C T C A G G C C A
Mutant sequence A G G C T C A G C C C T
ZBTB18
T A G C C A G A T G T G G
T A G C G T G A T G T G G
TFAP2C
WT sequence C A G C C T C A G G C C
Mutant sequence C A G G C T C A G C C C
ZNF384
C T A C A A A A A A A A
C T A C T A T T T A A A
MXI1
WT sequence G C C A C A T G G C
Mutant sequence G C C T C A A G G G
ZNF460
G C A T C A G C C T C C C G A G
G C A T C A G G G T C C C G A G
Bits


Figure S4

A

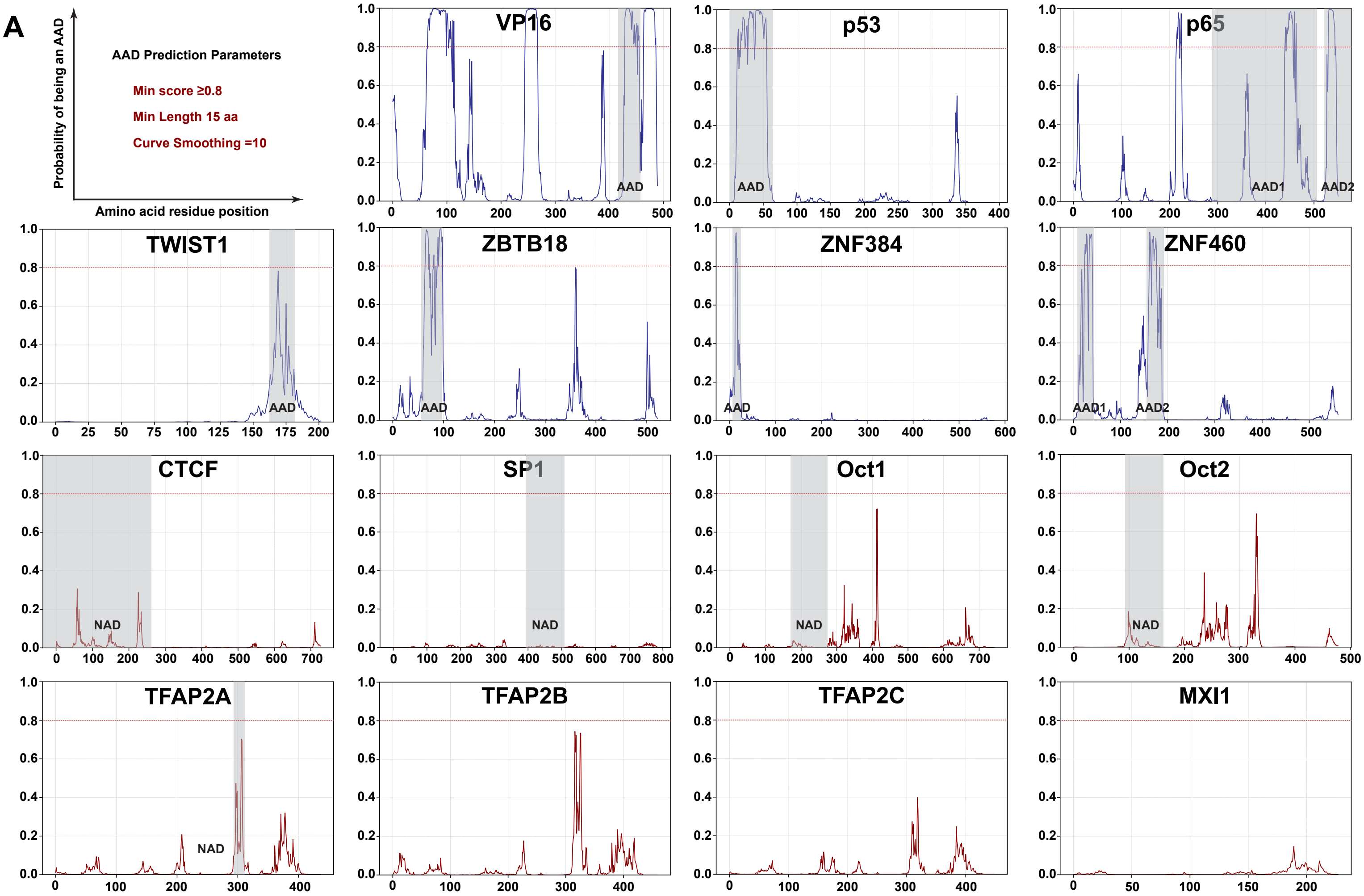


B

| Protein Name | Uniprot ID | Sequence of Tested AAD/NAAD | Amino Acid Position |
|---|---|---|---|
| VP16 | P06492 | APPTDVSLGDELHLDGEDVAMAHADALDDFDLDMLGDGDSP | 412-453 |
| p53 | P04637-1 | MEEPQSDPSVEPPLSQETFSDLWKLLPENNVLSPLPSQAMDDLMLSPDDIEQWFTE DPGPDEAPRMPEAAPPV | 1-73 |
| p65-AAD1 | Q04206-1 | FQYLPDTDDRHRIEEKRKRTYETFKSIMKKSPFSGPTDPRPPPRRIAVPSRSSASVP KPAPQPYPFTSSLSTINYDEFPTMVFPSGQISQASALAPAPPQVLPQAPAPAPAPAM VSALAQAPAPVPVLAPGPPQAVAPPAPKPTQAGEGTLSEALLQLQFDDEDLGALL GNSTDPAVFTDLASVDNSEFQQLLNQGIPVAPHTTEPMLMEYPEAITRLVTGAQRP PDPAPAPLGAP | 286-521 |
| p65-AAD2 | Q04206-1 | PGLPNGLLSGDEDFSSIADMDFSALLSQISS | 521-551 |
| DELQP | -- | DELQPASIDP | -- |
| TWIST1 | Q15672 | SDELDSKMASCSYVAHE | 165-181 |
| ZBTB18 | Q99592-1 | DIVHLNSDIVTAPAFALLLEFMYE | 61-84 |
| ZNF384-AAD | Q8TF68-1 | FWPSIPTVSGQIE | 12-24 |
| ZNF460-AAD1 | Q14592-1 | TFEDVAVTFTQEEWGQLDVTQRALYVEVM | 14-42 |
| ZNF460-AAD2 | Q14592-1 | GPVTDSLIHEGENSYKFEEMFNENCFLVQHEQ | 157-188 |
| CTCF NAAD | P49711 | MEGDAVEAIVEESETFIKGKERKTYQRRREGGQEEDACHLPQNQTDGGEVVQDVN SSVQMVMMEQLDPTLLQMKTEVMEGTVAPEAEAAVDDTQIITLQVVNMEEQPINIG ELQLVQVPVPVTVPVATTSVEELQGAYENEVSKEGLAESEPMICHTLPLPEGFQVV KVGANGEVETLEQGELPPQEDPSWQKDPDYQPPAKKTKKTKKSKLRYTEEGKDV DVSVYDFEEEQQEGLLSEVNAEKVVGNMKPPKPTKIKKKGVKKTFQCEL | 1-270 |
| SP1 NAAD | P08047 | ILIQPQLVQGGQALQALQAAPLSGQTFTTQAISQETLQNLQLQAVPNTGPIIIRTPTV GPNGQVSWQTLQLQNLQVQNPQAQTITLAPMQGVSLGQTSSSNTTL | 398-502 |
| Oct1 NAAD | P14859-1 | DLQQLQQLQQQNLNLQQFVLVHPTTNLQPAQFIISQTPQGQQGLLQAQNLLTQLP QQSQANLLQSQPSITLTSQPATPTRTIAATPIQTLPQSQS | 175-269 |
| Oct2 NAAD | P09086-1 | LAGDIQQLLQLQQLVLVPGHHLQPPAQFLLPQAQQSQPGLLPTPNLFQLPQQTQG ALLTSQPR | 98-161 |
| TFAP2A NAAD | P05549-1 | EGEAVHLARDFGYVCETE | 294-311 |

Figure S5
